# Transendothelial migration of the Lyme disease spirochete involves spirochete internalization as an intermediate step through a transcellular pathway that involves Cdc42 and Rac1

**DOI:** 10.1101/2024.09.10.612329

**Authors:** Daiana Alvarez-Olmedo, Claire Kamaliddin, Theodore B. Verhey, May Ho, Rebekah DeVinney, George Chaconas

## Abstract

Despite its importance in pathogenesis, the hematogenous dissemination pathway of *B. burgdorferi* is still largely uncharacterized. To probe the molecular details of transendothelial migration more easily, we studied this process using cultured primary or telomerase-immortalized human microvascular endothelial cells in a medium that maintains both the human cells and the spirochetes. In *B. burgdorferi* infected monolayers we observed ∼55% of wild-type spirochetes crossing the monolayer. Microscopic characterization revealed entrance points across the cellular surface rather than at cellular junctions, supporting a transcellular route. In support of this pathway, locking the endothelial junctions using a VE-PTP inhibitor did not reduce transendothelial migration. We also used inhibitors to block the most common endocytic pathways to elucidate effectors that might be involved in *B. burgdorferi* uptake and/or transmigration. Directly inhibiting Cdc42 reduced spirochete transmigration by impeding internalization. However, blocking Rac1 alone dramatically reduced transmigration and resulted in a concomitant increase in spirochete accumulation in the cell. Our combined results support that *B. burgdorferi* internalization is an intermediate step in the transendothelial migration process which requires both Cdc42 and Rac1; Cdc42 is needed for spirochete internalization while Rac1 is required for cellular egress. These are the first two host proteins implicated in *B. burgdorferi* transmigration across endothelial cells.

**IMPORTANCE:** Lyme borreliosis is caused by *Borrelia burgdorferi* and related bacteria. It is the most common tick-transmitted illness in the Northern Hemisphere. The ability of this pathogen to spread to a wide variety of locations results in a diverse set of clinical manisfestations, yet little is known regarding vascular escape of the spirochete, an important pathway for dissemination. Our current work has studied the traversal of *B. burgdorferi* across a monolayer of microvascular endothelial cells grown in culture. We show that this occurs by passage of the spirochetes directly through these cells rather than at cellular junctions and that internalization of *B. burgdorferi* is an intermediate step in the transmigration process. We also identify the first two host proteins, Cdc42 and Rac1, that are used by the spirochetes to promote traversal of the cellular monolayer. Our new experimental system also provides a new avenue for further studies of this important process.

## INTRODUCTION

Lyme borreliosis is a multi-system disease caused by *Borrelia burgdorferi* and related species (Coburn et al., 2021; Stanek and Strle, 2018; Steere et al., 2016). The pathogenic spirochetes are transmitted by the bite of an infected tick. After establishing a localized dermal infection, the spirochetes can disseminate via direct passage through soft tissue, lymphatic or circulatory systems. The latter is an important thoroughfare for dissemination to distal sites from the point of entry (Steere et al., 2016; Wormser, 2006; Wormser et al., 2008), making transmigration across the endothelium a crucial step in this pathway. Despite its importance, the hematogenous dissemination process remains largely uncharacterized, and the mechanisms involved in host-pathogen interactions that mediate spirochete extravasation are poorly understood. Remarkably, there are few reports regarding the mechanisms involved in *B. burgdorferi* transmigration through endothelial cells. A few early studies presented contradictory findings on *B. burgdorferi* extravasation, suggesting evidence for a transcellular and a paracellular route (Comstock and Thomas, 1989, 1991; Szczepanski et al., 1990). More recently, using intravital microscopy, we reported spirochete extravasation in living mice through a transcellular pathway in the microvasculature of the knee joint peripheral tissue (Tan et al., 2023).

Although most pathophysiological processes occur in the microvasculature, *in vitro* research on *B. burgdorferi*-endothelial interactions has predominately utilized HUVEC cells, which are macrovascular endothelial cells (Lafrance et al., 2011; Ristow et al., 2015; Thomas and Comstock, 1989). At present, only a few primary microendothelial cell culture systems have been reported that support spirochete transendothelial migration and the efficiency was low or not determined (see **Table S1**); in part, this may be due to a disparity in culture conditions supporting healthy growth of the spirochetes versus endothelial cells in culture.

In 1983, Barbour-Stoenner-Kelly (BSK) medium was developed, allowing the growth of *B. burgdorferi* in laboratory settings (Barbour, 1986). In particular, this medium is rich in lipids and contains essential nutrients for the spirochetes, such as cholesterol, adenine, spermidine, and N-acetyl-glucosamine (Barbour, 1986; von Lackum and Stevenson, 2005; Wyss and Ermert, 1996). However, BSK is unsuitable for eukaryotic cells, especially primary endotheial cells, which have specific requirements for growth, such as hydrocortisone and growth factors (vascular and endothelial growth factor (VEGF/ EGF)(Ferrara et al., 1992). Moreover, any conditions that might activate the endothelial cell monolayer lead to cell-cell junction opening, altering endothelial integrity and the corresponding barrier function (Wyss and Ermert, 1996). Consequently, it has been challenging to develop a co-culture medium that satisfies the needs of both the host endothelial cells and *B. burgdorferi*. Few studies have been published regarding *B. burgdorferi’s* interaction with various primary and endothelial cells (see **Table S1** for details). Among these studies and others, different media compositions were tested, but none maintained optimal endothelial cell and spirochete viability.

To overcome this obstacle and to study the interaction between *B. burgdorferi* and human endothelial cells *in vitro*, we developed co-culture conditions using a medium suitable for both participants. We were particularly vigilant towards *B. burgdorferi* and endothelial-cell morphology, growth, and cytotoxicity. Moreover, in our *in vitro* model, *B. burgdorferi* and human primary endothelial cells happily coexist with a stable cellular monolayer for up to 48 hours and support efficient transendothelial migration to about 55% of input spirochetes. Here, we use these co-culture conditions to investigate spirochete penetration and endothelial migration using microscopic analysis and chemical inhibitors of the most common endocytosis pathways. Our findings support a transcellular pathway for *B. burgdorferi* extravasation that uses cellular facilitation by Cdc42 and Rac1, members of the Rho family of GTPases.

## MATERIALS AND METHODS

### Bacterial strains and culture

Spirochete cultures in BSK-II medium (Barbour, 1984) containing 6% rabbit serum (31125, Pel-Freez) were inoculated from frozen glycerol stocks. The spirochete cultures were grown for 48 hours at 35°C to a concentration of 1-5 x 10^7^ before infecting the cells. The predominant *B. burgdorferi* strains used (Moriarty et al., 2008) were GCB726: low passage, infectious, GFP-expressing B31 5A4-NP1 (Kawabata et al., 2004) and GCB705: high passage, non-infectious, non-adherent, GFP-expressing B31-A (Bono et al., 2000). These strains were grown in media containing 100 μg/ml gentamycin (4730, Omnipur, Calbiochem). For details on all strains used, see **Table S2**.

### Human cell culture

Primary human dermal microvascular endothelial cells (hMVEC-d) were purchased from Lonza (CC-2543), grown in EBM (CC-3156, Lonza) complete media (with supplements EGM^TM^-2 SingleQuots^TM^ Supplements (CC-4176, Lonza)) at 37°C under 5% CO2, and used before passage five. hTERT-immortalized Dermal Microvascular Endothelial Cell, Neonatal (CRL4060, ATCC), was used. hTERT was cultured in Basal medium (PCS-100-030, ATCC) with the Microvascular Endothelial Cell Growth kit-BBE (PCS-110-040, ATCC) +0.5 ug/ml puromycin (P8833, Sigma-Aldrich) according to the manufacturer’s instruction.

### Inhibitors

When indicated the following inhibitors were used: LY294002 (S1105, Selleckchem), U0123 (9903, Cell Signaling), cilengitide trifluoroacetate (S7077, Selleckchem), filipin III from *Streptomyces filipinensis* (480-49-9 Sigma-Aldrich), dynasore (304448-55-3, EMD millipore), desatinib (BMS-354825, S1021, Selleckchem), 2-Fluoro-N-[2-(2-methyl-1H-indol-3-yl)ethyl]-benzamide ((CK-666), SML0006, Sigma-Aldrich), imipramine hydrochloride (I0899, Sigma-Aldrich), SB203580, 5-(N-Ethyl-N-isopropyl) amiloride (1154-25-2, Sigma-Aldrich), ML141 (71203-35-5, EMD millipore), EHop-016 (S7319, Selleckchem) and NSC 23766 trihydrochloride (S8031, Selleckchem).

### Co-culture

We developed the co-culture (CoMe) with the objective of maintaining the microendothelial cells and the *B. burgdorferi* during suitable times for infections. For preparing the CoMe, a modified BSK (BSK-/-) lacking rabbit serum and HEPES was used in conjunction with M199 (31100035, Gibco) prepared as recommended by the manufacturer and buffered with Na_2_CO_3_ (S5761, Sigma) and 10% human serum (HS, pooled Human AB Serum plasma-derived ISERAB100ml, Innovative Research Inc, cat 50-203-6404). Note that the M199 medium has sodium bicarbonate as a buffer, so HEPES was excluded from the CoMe recipe; moreover, instead of using rabbit serum, which was detrimental for the human cells, we used human serum. Different mixes of co-culture medium were prepared using various proportions of BSK-/- plus M199 and 10% HS: 1/3 M199+2/3 BSK (-/-) (final CoME preparation) or ½ M199+1/2 BSK (-/-). The final CoMe mix was filtered sterile using a 0.22 μm filter and supplemented with 10 ng/ml EGF (CC-4107, Lonza) and 1 ug/ml hydrocortisone (H0888, Sigma). This CoMe was stored at 4°C for up to four weeks and never frozen to avoid crystal formation.

### Human cell morphology assessment

Cell morphology was assessed by using fluorescence microscopy. Briefly, cells were seeded on borosilicate glass round coverslips (12 mm diameter, 0.13-0.17 μm thickness, Fisher Scientific) coated with 300 µl 10% rat tail collagen type I (354236, Corning; 3.4 mg/ml) in 0.2M acetic acid for 1 h (Corning, cat 354236). After incubation in the corresponding media, cells were washed with warm HBSS, and fixed for 10 min in 2% paraformaldehyde (EMS). Permeabilization was carried out with 0.5% Triton X-100 and labelling with either 1/200 VE-cadherin conjugated Alexa Fluor 647 (A22287, Invitrogen™, 1/200) or Alexa Fluor 546 Phalloidin (A22283, Invitrogen™ 1/200) for F-actin. *B. burgdorferi* was labelled with primary rabbit antibody against *B. burgdorferi* (1439-9406, Bio-Rad, 1/100), followed by anti-rabbit Cy3 labelled second antibody (Molecular Probes, 1/200). All incubations were performed at room temperature. Coverslips were mounted using ProLong™ Gold Antifade with DAPI (P36931, Invitrogen) and imaged using a Leica DMIRE2 wide-field microscope with the 40X oil immersion objective. Acquired micrographs were analyzed using Volocity Version 6.5.1 (Perkin Elmer).

### Media validation: *B. burgdorferi* growth curve and morphology analysis

To ensure *B. burgdorferi* well-being in the co-culture media, we performed growth assays in different conditions. 3x10^5^ spirochetes were seeded in the indicated media and enumerated at 24 h intervals over 48 h by darkfield microscopy (20x, Nikon eclipse E400) using a Petroff Hauser chamber. After 48 hours, spirochetes were motile in all assayed media except the EBM.

Since *B. burgdorferi* can be filamentous in non-optimal culture conditions, we specifically assessed spirochete length in the tested media (1/3 M199+ 2/3 modified BSK-II+10% HS or ½ M199+ 1/2 modified BSK-II+10% HS) compared to the regular BSK-II by fluorescence microscopy. Briefly, GCB726 was grown for 48 h in either BSK-II or the indicated media and images were acquired with a 40X oil immersion objective on a wide-field Leica DMIRE2 microscope (Leica, Wetzlar, Germany). Images were acquired using an ORCA-ER digital camera controlled with Openlab (Improvision, Coventry, UK) software. Spirochete length was assessed using Fiji for ImageJ (64-bit) 2.0.0 (Schindelin et al., 2012). The length of 60 spirochetes was analyzed in each media condition using the Freehand line tool in Fiji. The length data were analyzed and plotted using GraphPad Prism version 8.0.0 for Windows.

### Viability

Cell metabolic activity was determined using water-soluble and cell-permeable 2-(4-iodophenyl)-3-(4-nitrophenyl)-5-(2,4-disulfophenyl)-2H-tetrazolium monosodium salt (WST-1), (5015944001, CellPro-Ro Roche) reagent according to the manufacturer’s instructions. Briefly, the nonradioactive stable tetrazolium salt WST-1 is cleaved to a soluble formazan when NAD(P)H is available in viable cells. Therefore, the amount of formazan dye formed directly correlates to the number of metabolically active cells in the culture. For the experiment, cells were seeded in a 96-well-plate until confluence was reached in either EBM-2 (for hMVEC-d) or VCBM (for hTERT) and then changed to EBM-2 or CoMe for 24 or 48 h. Next, 10 µl was added to each well and incubated for 2 h, before reading using a Spectramax i3x plate reader at 440 nm. In all the cases, each plate contained blanks, controls, and the respective treatments; three independent experiments were performed at least in triplicate (n≥9).

### Cytotoxicity Assay

Cytotoxicity was measured in hTERT with CytoTox96 (Promega) according to the manufacturer’s instructions. Briefly, this assay measures lactate dehydrogenase (LDH), a cytosolic enzyme released upon cell lysis. The enzyme converts a tetrazolium salt (iodonitrotetrazolium violet) into a red formazan product, and the color intensity is proportional to the number of lysed cells. The percent of cytotoxicity was calculated with the following equation: % cytotoxicity= 100*Experimental LDH Release (OD_490_)/Maximum LDH released (OD _490_). On each plate, the manufacturer-recommended controls (No-cell, vehicle-only cells and maximum LDH Release control) were performed. The Maximum LDH release was calculated by lysing the cells as indicated by the manufacturer.

### Confocal imaging

Stained samples were imaged on a Nikon AR1 multichannel confocal microscope (Nikon, Melville, NY, USA) in the Snyder Live Cell Imaging facility (LCI) at the University of Calgary. The microscope comprised of a Ti2 flagship inverted microscope paired with a Ti2 XY drive and a Ti2 Z drive with an A1 Piezo Z Drive fitted with a motorized objective turret. The scanner was set on Galvano mode (high-resolution Galvano scanning). Image acquisition was performed using Nikon NIS-Elements (AR v 5.02.00) software using either 40X oil immersion Plan Fluor DICN2 (working distance 0.24 mm 1.30 NA) for spirochete entry assay. For intracellular target analysis, image acquisition was performed with a 60X oil immersion objective. Laser excitation wavelengths of 403.3, 488.2, 561.9, and 638.6 nm were used in rapid succession. Laser power was adjusted in each experimental condition. The pinhole size was set up for a 638.6 nm wavelength.

### Transmission electron microscopy (TEM)

To evaluate the ultrastructure of the interaction between the spirochete and hMVEC cells, the microendothelial cells were plated on cell culture-treated sterile 25 mm diameter plastic coverslips (174985, Nunc Thermanox), placed in 6 well plates and grown to confluence in EBM (CC-3156, Lonza) complete media (with supplements EGMTM-2 SingleQuotsTM Supplements (CC-4176, Lonza) at 37°C under 5% CO2. The cells were infected with 8x10^7^spirochetes in CoMe for 20 h. After the incubation, the cells were fixed with glutaraldehyde 2.5 % buffered to pH 7.4 in 0.1M sodium cacodylate buffer for a minimum of 2 hours. The specimens were washed in 0.1 M sodium cacodylate buffer at pH 7.4 before being post-fixed in 2% osmium tetroxide. The samples were dehydrated with graded acetone, infiltrated with several changes of graded Epon: Acetone, and then embedded in Epon resin. The sections were cut at 70nm, stained with a 2% uranyl acetate and counterstained with a 4% lead citrate solution. The TEM images are taken on a Hitachi model H-7650 with an AMT16000 camera.

### Transmigration analysis

Cells (hMVEC-d or hTERT) were seeded in the upper chamber of the Transwells (10769-240, VWR insert, 24 wells, PET membrane, 3.0 µm) at a density of 3 × 10^5^ in 100 µl of media; in the lower chamber 650 µl of media was added. The cells were grown to confluence (2-3 days) without puromycin for hTERT. The confluence was confirmed in at least one of the tested Transwells before each experiment using an albumin diffusion assay (see below). After the infection, the total media in the upper and lower chamber was analyzed for spirochetes, either by microscopic counting using a Petroff-Hauser chamber or by flow cytometry.

### Flow cytometry

For transmigration calculations, the medium in the upper (100 µl) and lower (650 µl) chambers was collected after a 20 h infection. The samples were washed twice by centrifugation in Eppendorf tubes at 6000 *× g* at 4**°**C in a tabletop centrifuge for 15 min, followed by resuspension in 500 µl of sterile filtered phosphate-buffered saline (PBS) (NaCl 0.137M, KCl 0.0027M, Na_2_PO_4_ 0.01M, KH_2_PO_4_ 0.0018M, pH=7.4). The washed spirochetes were transferred to 5 ml analysis tubes (Polystyrene tubes). The sample was analyzed with the BD FACSCanto 3-laser instrument. The cytometer was set with a threshold of FSC 400 and SSC 200 for visualizing small cells or bacteria. The PBS was run first at high, medium, and low speeds for 1 min each to determine the noise level. Subsequently, the rest of the experiment was performed at low speed. A negative control, *B. burgdorferi* without GFP, was used to set the cutoff. A positive Counts versus FITC-A (GFP) control and the samples were also run. The percentage of transmigration was calculated as the % bottom/[top + bottom] of *B. burgdorferi* recorded in one minute. For *B. burgdorferi* internalization, hTERT cells were grown in T-25 flasks to confluence (1.5x10^6^ total cells) and infected with *B. burgdorferi* at a multiplicity of 7:1. After 20 h, the cells were treated with trypsin, washed with HBSS, fixed with PFA at 0.25%, and run on the flow cytometer. A hundred thousand cells were analyzed, and the percentage of GFP-infected cells and the median fluorescence intensity (MFI) were quantified. MFI was used for comparisons btween the different treatments (infection or infection+amiloride)

### Albumin diffusion assay

Transendothelial electric resistance (TEER) levels across microvascular endothelial cells is very low and not a sensitive enough measure to assess changes in monolayer integrity. Therefore, we have used the more sensitive approach of measuring albumin permeability (Bednarek, 2022). Monolayer integrity was measured by adding 10 µg of bovine serum albumin conjugated to Alexa Fluor 555 (Thermo Fisher Scientific, cat A34786) in the upper chamber. After 4 h, the media in the upper and lower chambers was collected for analysis. The optical density was measured with a fluorescence spectrophotometer (SpectraMax i3x Multi-Mode Microplate Reader) at wavelengths of 525 nm excitation and 560 nm emission. The percentage of diffusion = lower chamber fluorescence*100/ (upper chamber fluorescence + (6.5*lower chamber fluorescence)). The coefficient 6.5 is the relationship between the volume in the upper and lower chambers.

### Statistics and data analysis

Analysis of micrographs was performed using ImageJ 1.53a software (Rasband, W.S., ImageJ, U. S. National Institutes of Health, Bethesda, Maryland, USA, https://imagej.nih.gov/ij/, 1997-2018). Statistical analyses were completed using GraphPad Prism version 9.5.0 for Windows, GraphPad Software, San Diego, California USA, www.graphpad.com. A non-parametric t-test or non-parametric ANOVA with the indicated post-test was used and indicated in the legend of each figure.

### Analysis of penetration point coordinates and endothelial cell junctions

A custom Python script for analyzing the shortest distance to the perimeter from the penetration point that accounts for the proportion of the cell surface area that is considered close to the junction was used and is available at https://github.com/verheytb/JunctionDistance. Briefly, a single-cell analysis in 18 cells that displayed clear staining in a single field of view was performed. First, for each cell, the perimeter and penetration point coordinates were measured using Fiji image analysis software. Then, with the script, we calculated the probability that the penetration point was closer to the nearest edge in two-dimensional space than expected by chance by using a null distribution of 10,000 points generated from a continuous uniform distribution within the cell’s perimeter. We measured the shortest distance to the perimeter for the penetration point (*d_pp_*), and for all randomly generated points *N* = {*d*_*r*1_ ⋯ *d_rn_* }, and then normalized the distance of the penetration point using the null distance distribution such that 0 < *n_pp_*< 1:

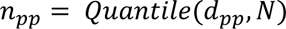

The normalized distance, *n_pp_*, enables comparison between cells of multiple cell sizes and shapes. A quantile of 0 represents an entry point at the endothelial junction, 0.5 is the median distance to the boundary, and 1 represents an entry point as far away from the boundary as possible.

## RESULTS

### Validation of the Co-Medium for *B. burgdorferi* maintenance

With the objective of generateing a CoMe that sustained *B. burgdorferi* and endothelial metabolic activity during a suitable time for infections, we tested diferent media combinations. We started by individually adding the following media to BSK-II (without rabbit serum and HEPES) at different ratio concentrations: M199, DMEM, RPMI, Cell Systems Endothelial Cell Medium, and DMEM: Ham’s F12. Only M199 showed potential to be part of the co-culture medium, and it was added to BSK-II without rabbit serum and HEPES in proportions of 1/3 or 1/2; the rest of the media tested were not used for further experimentation because they induced filamentous (50 µm and longer) spirochetes, death, and formation of “rosettes” or aggregates (data not shown). Other authors have also shown some potential of M199 for short-term *B. burgdorferi in-vitro* experiments (see **Table S1**). As shown in **Fig. 1A**, *B. burgdorferi* growth was assayed either in complete BSK-II (used as positive control), EBM-2 (complete cell medium, used as negative control), or a mix of modified BSK-II and M199. The mixes were either 1:3 (M199:BSK-II), which we refer to hereafter as CoMe, or 1:2 (M199:BSK-II). For these mixes, a modified BSK-II was used, lacking HEPES (because the M199 medium was already buffered with sodium bicarbonate) and rabbit serum (because we found it was detrimental for cell growth), and supplementing instead with 10% human serum to fulfill the spirochete and cell requirements. The CoMe conditions showed no difference in growth rate when compared with the control BSK-II complete medium and, therefore, was a good medium to maintain spirochete growth for up to 48 h. Next, we investigated whether CoMe induced any morphological changes; however, no formation of rosettes or aggregates was visualized. The length of the spirochetes was measured after 48 h, and no differences were observed between the CoMe and the complete BSK medium (**Fig. 1B**).

**Fig. 1.**
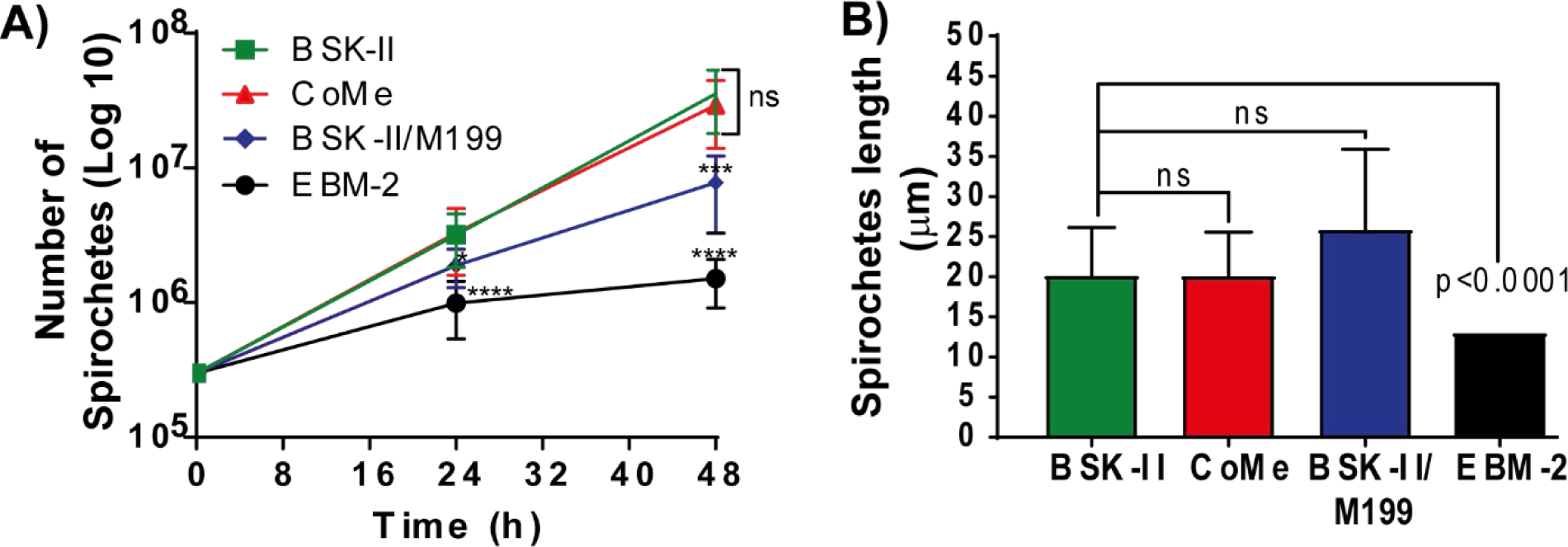
Validation of the CoMe for *B. burgdorferi*. **A)** Evaluation of B. *burgdorferi* growth rate in the indicated media: Infectious *B. burgdorferi* spirochetes (GCB726) were grown in BSK-II. Using log phase spirochetes, each of the studied media was inoculated with 3x10^5^ spirochetes/ml: CoMe (red triangle; optimized co-culture media), BSK (green square; *B. burgdorferi* reference media), EBM (blue circle; cell reference media), and 1:1 BSK-II/M199 supplemented with 10% human serum (black diamonds). Spirochetes were enumerated every 24 h using a Petroff-Hauser chamber. The graph represents at least three independent experiments; the y-axis is shown in log_10_ scale. The error bars denote standard deviation, and significance was investigated using two-way ANOVA with mixed effects with Geisser-Greenhouse correction and Dunnett’s multiple comparison test. **B)** Evaluation of the length of *B. burgdorferi* in different media. After 48h of incubation in the desired medium, the spirochetes were imaged using a LEICA DMIRE2 fluorescent microscope at 400x magnification. The length of 60 spirochetes per condition was assessed using the freehand line tool in Fiji. The graph represents the values +SD from 3 independent experiments; *p* <0.05 was considered significant.

### Validation of the Co-Medium for endothelial cell maintenance

In general, endothelial cells require specific culture media accompanied by several growth factors to stimulate endothelial cell proliferation without affecting endothelial phenotype or function (Leopold et al., 2019). The objective of developing the CoMe was not to induce growth in cells but to provide the essential components that allow the maintenance of optimal metabolic activity and monolayer in the cells and would also support spirochetes for up to 48 h. This would allow studying the interactions between *B. burgdorferi* and the endothelial monolayer during the transmigration process. Therefore, to evaluate if CoMe would suit our objective, two endothelial cells were selected: hMVEC-d, which are primary dermal endothelial cells from a female donor, and hTERT, immortalized dermal endothelial cells from a male donor. Cells were grown to confluence in (EBM-2 or VCBM as indicated), then the media was exchanged for CoME, and finally, the metabolic activity was measured using WST-1. No significant differences were found between the tested cell medium and the proposed CoMe (**Fig. 2A**). We also analyzed whether the CoMe was cytotoxic to hMVEC-d cells using a tetrazolium salt and assessing the percentage of cytotoxicity at times varying from 15 to 50 h. The CoMe showed a significantly lower cytotoxicity compared to EBM-2 (**Fig. 2B**). These results together suggest that CoMe is a good medium to safely use to maintain endothelial cells and *B. burgdorferi* for periods up to 48 h.

**Fig. 2.**
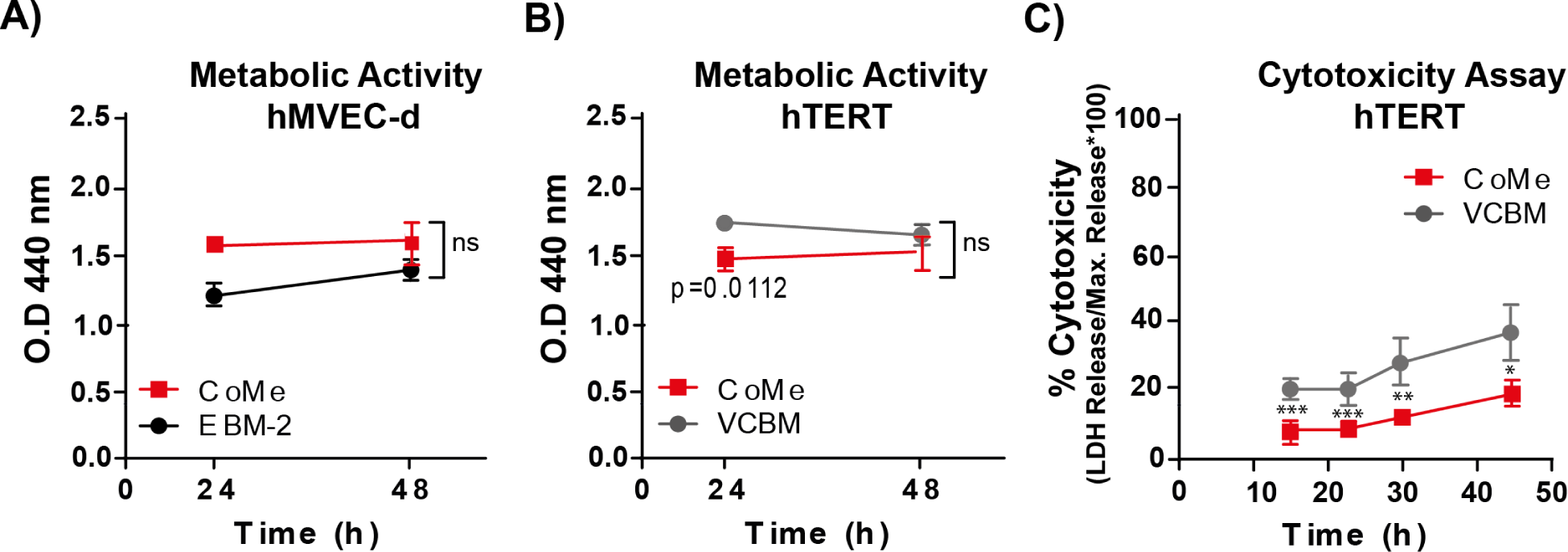
Validation of the CoMe in hMVEC-d and hTERT cells. **A, B)** The metabolic activity of hMVEC-d **(A)** and hTERT **(B)** cells was evaluated by measuring the metabolic activity with the WTS-1 reagent, a cell-permeable tetrazolium salt. The cells were seeded in 96 well plates and grown until confluent in EBM-2 (I) or VCBM (II). The medium was then changed to EBM-2 or CoMe at 24 and 48 h. The amount of formazan released was measured at 440 nm. The graph represents the results of three independent experiments performed at least in triplicate (n>9) and analyzed with one-way ANOVA (Kruskal-Wallis test). No significant changes were observed between the reference media (VCBM) and CoMe after 48 h treatment. **C)** The kinetic analysis of cytotoxicity was evaluated by the release of lactate dehydrogenase (LDH). hTERT were seeded in 96 well plates and grown until confluent. The medium was then changed to EBM-2 or CoMe for 16, 32, 24, and 48 h. The graph represents the results of three independent experiments (n >10) analyzed with the Kruskal-Wallis test; *p* <0.05 was considered significant. The discontinuous blue line represents the basal cytotoxicity levels in MVEC cells in the studied medium.

### Analysis of *B. burgdorferi-*endothelial interactions

To evaluate the different types of interactions between *B. burgdorferi* and endothelial cells, a dual staining experiment was performed using GFP-expressing *B. burgdorferi*. The cells were then fixed and stained with anti-*B. burgdorferi* whole-cell antibody/Cy3 (pink) without permeabilization to label extracellular bacteria. This process allowed us to discriminate between: 1) adherent extracellular spirochetes (**Fig.3A** left panel, represented in pink), 2) penetrating spirochetes (**Fig. 3A**, middle panel, dual color, yellow arrow) and 3) intracellular spirochetes (**Fig. 3A**, right panel, represented in green). The analysis showed that of 4,302 spirochetes analyzed, 75% were adherent or extracellular, 25% intracellular, and only 1% penetrating (**Fig. 3B**).

**Fig. 3.**
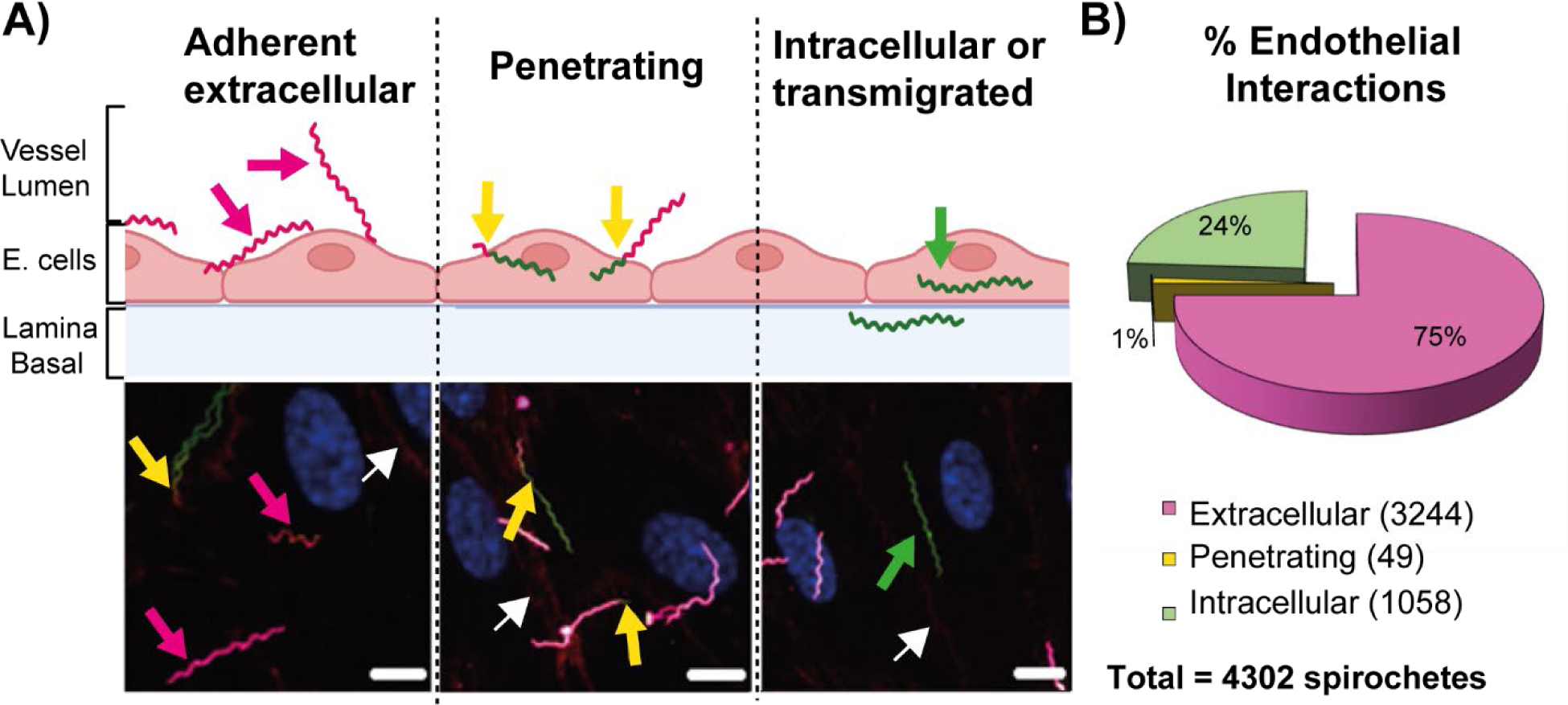
Analysis of endothelial interactions: adhesion, penetration, and internalization. **A)** Scheme of the three endothelial interactions evaluated (upper panel) and a corresponding representative micrograph for each (lower panel) obtained by confocal microscopy. In all cases, extracellular (adherent) spirochetes appear in pink (see pink arrow left panel), intracellular spirochetes in green (see the green arrow, central panel), and penetrating spirochetes are bicolor (see yellow arrows). Scale bar = 6 μm. For the micrographs, hMVEC-d seeded on collagen-coated coverslips were infected with GCB726, fixed, washed, and labelled as noted in Materials and Methods. Spirochetes adhered to the extracellular surface of the cell were discriminated from the intracellular ones by labelling with rabbit anti-*B. burgdorferi* antibody followed by an anti-rabbit secondary antibody (Cy3). Conjugated VE-cadherin (AlexaFluor 667) labelling (red, indicated by white arrows). **B)** Analysis of *B. burgdorferi*-endothelial interactions: a total of 4,302 spirochetes were analyzed at 16 hours post-infection by counting in four independent experiments. Among these spirochetes, 3,244 (75%) were extracellular, 1,058 (24%) were intracellular or transmigrated, and 49 (1%) were penetrating.

### Analysis of adherent spirochetes

The external spirochetes expressing GFP were labelled with whole-cell *B. burgdorferi* antibody (Cy3) to differentiate them from internalized spirochetes (**Fig. 3A**), left panel. Approximately 31% of the adherent spirochetes were at cellular junctions, and 69% were found distal to the junctions (**Fig. 4**). These results suggest that under the conditions of this experiment spirochete adherence takes place preferably on the cellular surface. However, the precise distribution cannot be determined from this experiment because the surface area corresponding to junction versus cell is not possible to calculate and the spirochetes themselves are 10-30 µm and do not adhere to a single defined location. To overcome these problems, we analyzed the penetration points of spirochetes that can be localized according to specific cellular positions and their distance from the cell boundary as determined by a custom script (see next section and **Fig. 5**).

**Fig. 4.**
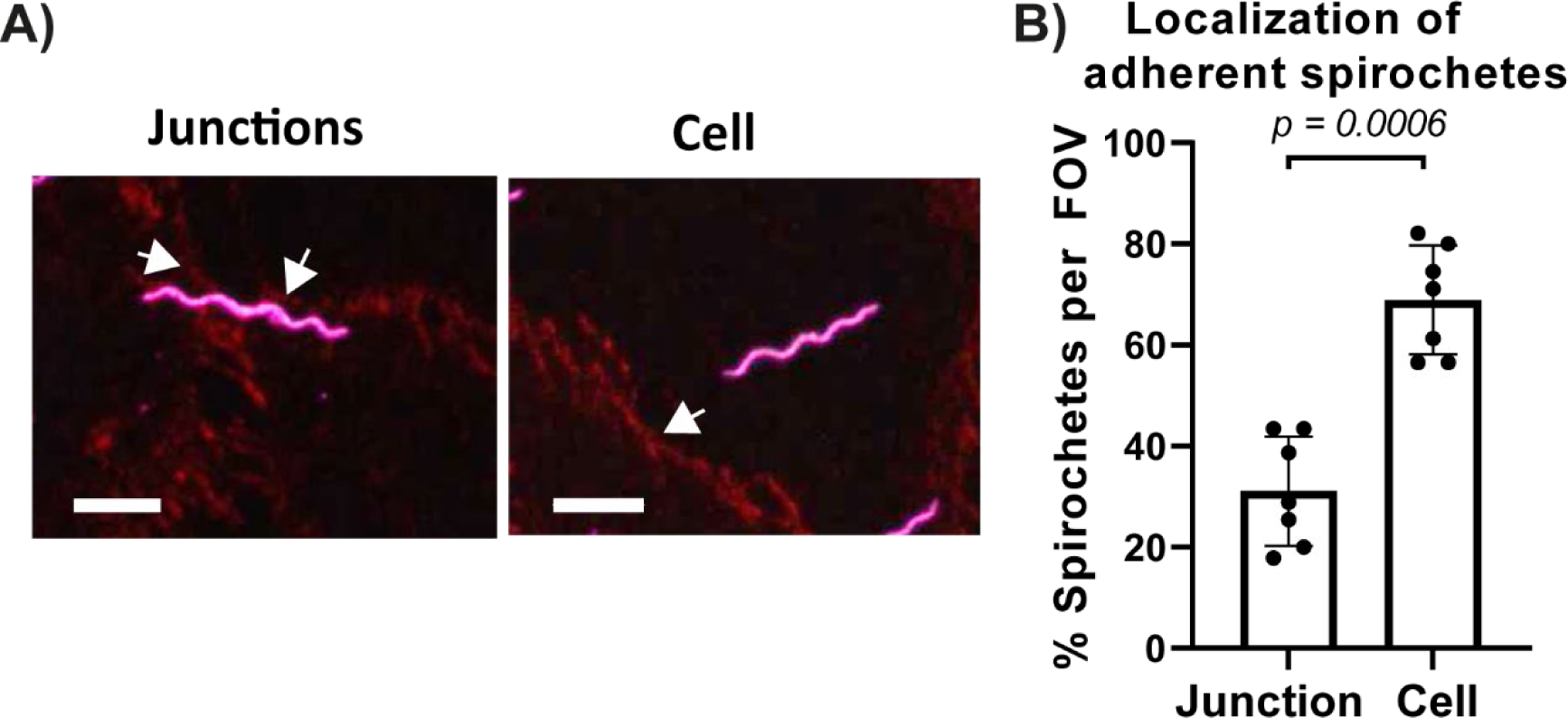
Analysis of endothelial adhesion of *B. burgdorferi*. **A)** Analysis of adhesion of *B. burgdorferi*: the micrographs are projection views of a z-series (extended focus) obtained by confocal microscopy. hMVEC-d cells seeded on collagen-coated coverslips and infected with *B. burgdorferi* GCB726 (WT) expressing GFP for 16 h in CoMe. The cells were washed thoroughly, fixed, and stained with VE-Cad conjugated to AlexaFluor 647 (junctions delimited with VE-cadherin are shown red in the figure, and are indicated by white arrows) and antibody anti-*Bb*/Cy3. *B. burgdorferi* GCB726 is the wild-type strain positive for anti-*B. burgdorferi*/Cy3 and displayed in magenta in the figure, scale bar = 6 μm. **B)** The graph represents the percentage of total adherent *B. burgdorferi* located either at the cellular junctions or on the cell surface. The position of 439 extracellular spirochetes (GCB726) adhered to HMVEC-d cells was assessed. Localization was analyzed successfully for 335 spirochetes. 31.1 ± 10.8 % of spirochetes were adherent to the junction, and 68.9 ± 10.8 % were adherent to the cellular surface. The data represents the percentage +/- SD of spirochetes at a given location per field of view (FOV) (n = 7 FOV); *p*-values < 0.05 were considered significant. Statistical analysis was carried out with the Mann-Whitney test, *p* = 0.006.

**Fig. 5.**
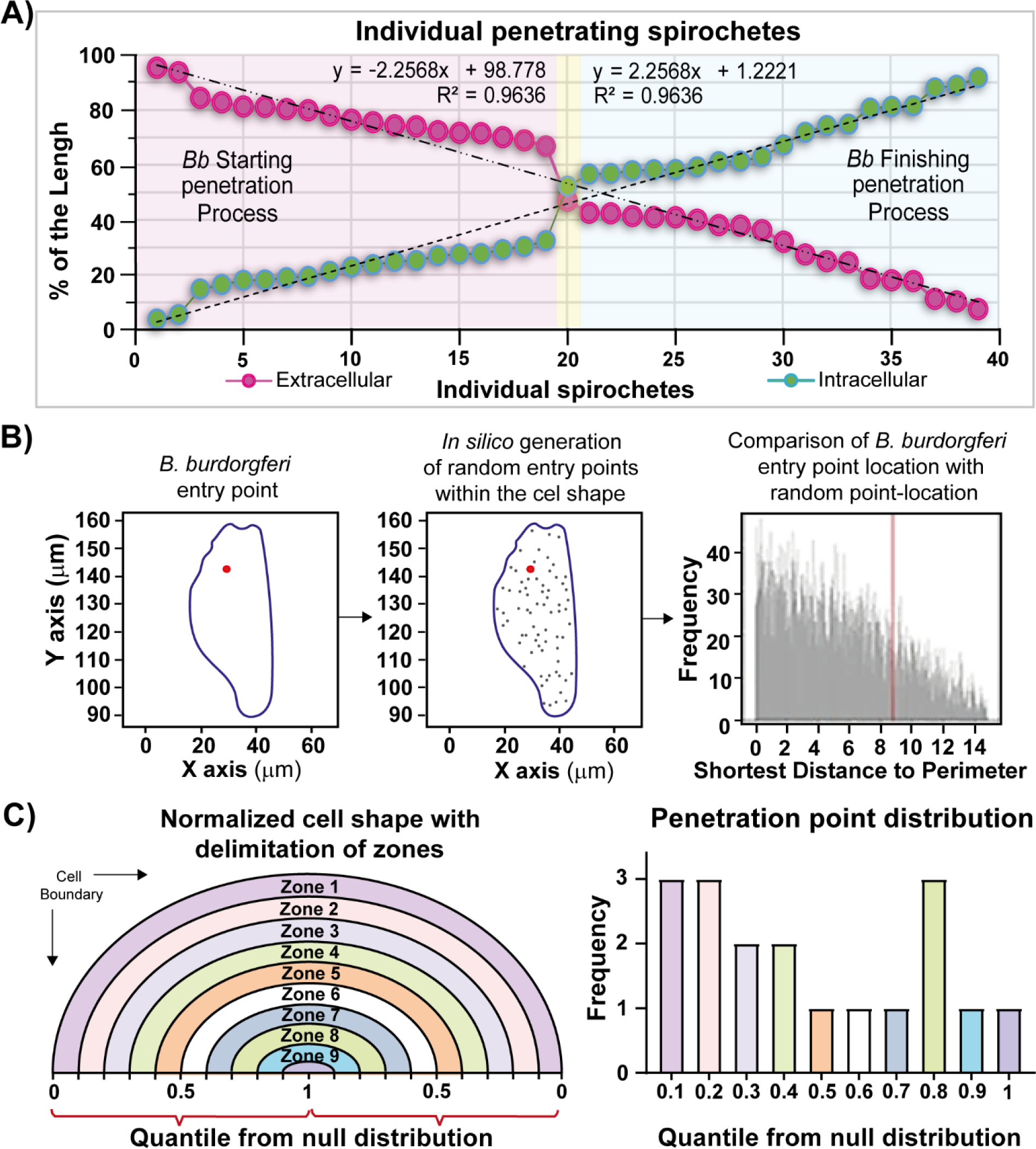
Analysis of endothelial penetration by *B. burgdorferi*. **A)** Evaluation of each single penetrating *B. burgdorferi* (strain GCB726): spirochete length is represented as the percentage of the length corresponding to the intracellular portion of *B. burgdorferi* (green dots) and the % length of the extracellular portion (pink dots). **B)** Example of the analysis methods used to analyze each cell containing a penetrating B. *burgdorferi*: micrographs showing penetrating spirochetes were used to identify coordinates of the perimeter (blue) and penetration point (red). The perimeter was used to generate randomly distributed points from a continuous uniform distribution (middle panel). Finally, the shortest distance to the perimeter was calculated for the penetration point (red) and the null distribution (grey). **C)** Analysis of the location of each entry point relative to the population. The illustration on the left is a schematic of the quantile distribution normalized to an ideal cell shape. Note the location of the zones analyzed (zones 1-10), where each zone is 10 % of the distance to the closest boundary. The graph on the right shows the frequency of the distribution of the penetration points in each quantile from all the penetration points analyzed (n= 18 cells). The quantile = 0 represents points closer to the boundary than all simulated points; 0.5 represents the median distance, and 1 signifies the points furthest away from the edge compared to simulated points.

### Characterization of *B. burgdorferi* penetration of endothelial cells

To analyze the penetration step in *B. burgdorferi* transmigration (**Fig. 3A**, right panel), we performed a single-cell analysis, considering each penetration point of *B. burgdorferi*. First, to ensure we had representative stages of the penetration process, we determined the extracellular/intracellular percentage of the individual penetrating spirochetes (**Fig. 5A**). There was a wide distribution of the observed penetration process, with a linear correlation coefficient of R^2^= 0.9636. Half of the evaluated spirochetes were initiating the penetration process (less than 45% of the spirochete length was found to be intracellular); the other half were finishing the process (more than 55 % of the *B. burgdorferi* length was located intracellularly), and one was in the middle of the process (between 46 and 54 % intracellular). This suggests that penetration is a continuous process that may occur at a uniform speed from the onset through its conclusion.

Early literature was exceedingly limited but controversial as to whether vascular transmigration was transcellular (Comstock and Thomas, 1989, 1991; Szczepanski et al., 1990) or paracellular (Moriarty et al., 2008; Szczepanski et al., 1990). However, more recent experiments suggest that in the mouse knee joint, a transcellular pathway of vascular transmigration is used (Tan et al., 2021). To further study the penetration process, we performed a single-cell analysis, which offers the potential to test the spatial co-localization of the penetration point with the cellular boundaries. Based on imaging data, a simple metric for assessing this in individual cells is to analyze the distance from the penetration point to the nearest cellular boundary.

To analyze the population of cells with penetrating *B. burgdorferi*, we assessed whether the penetration point was associated with endothelial boundaries in 2D space while accounting for the diverse size and shape of cells. To do this, we used imaging data from 18 cells that displayed clear staining in a single field of view. Thirty-one cells undergoing penetration with *B. burgdorferi* were excluded because of incomplete membrane staining (either poorly imaged or part of the cell was outside the field of view during image acquisition). For each cell, perimeter and penetration point coordinates were measured using Fiji image analysis software (**Fig. S1**). These 2-dimensional coordinates were used as inputs to a custom script, which calculated the probability that the penetration point was closer to the perimeter than expected by chance in two-dimensional space. We normalized the distance of the penetration point with a null distribution of 10,000 points generated from a continuous uniform distribution within the perimeter of the cell to account for differences in cell shape and size (**Fig. 5B**).

Finally, the penetration point was evaluated as the shortest distance to the perimeter and was converted to a quantile from the null distribution to enable comparison between cells of multiple cell sizes and shapes. A quantile of 0 represents an entry point at the endothelial junction, 0.5 is the median distance, and 1 represents an entry point as far away from the boundary as possible. By doing this, we summarized the normalized distances into ten zones of an ideal cell and plotted the frequency for each zone (**Fig. 5C**). Although there were a limited number of cells used for the analysis (n= 18 cells), we found at least one *B. burgdorferi* entry point in each of the 10 zones of the distribution, suggesting that the penetration process can occur across the surface of the cell rather than specifically near endothelial junctions.

### Analysis of internalized *B. burgdorferi*

It is known that intracellular spirochetes can be found as debris or as circular or elongated forms (Wu et al., 2011). Circular spirochetes are considered stressed, a form that *B. burgdorferi* can adopt in the presence of antibiotics or to avoid immune clearance (Wu et al., 2011). All the intracellular spirochetes observed (n=1058, 24% of total) showed an elongated shape (**Fig. 3A** right panel), consistent with transmigrating spirochetes, not with stressed or degrading forms. Previous findings in non-phagocytic cells indicate a wide range of percentages (from 10-25%) of internalized *B. burgdorferi,* which seems to depend on the cell type studied and perhaps the presence of growth factors (Ma et al., 1991). Further details are noted in the Discussion.

### Assessment of the transendothelial migration pathway using Transwell chambers

To move beyond the internalization of *B. burgdorferi,* we used Transwell chambers to monitor complete endothelial transmigration by the spirochetes (Nyarko et al., 2006; Van Gundy et al., 2021). Using our CoMe, we established an efficient endothelial transmigration system (**Fig. 6A**, see Materials and Methods) using hMVEC-d (human primary dermal (female donor); left panel) and human hTERT (male telomerase-immortalized dermal; right panel) endothelial cells. Before infection at an MOI of 7, the monolayer integrity was verified by diffusion of less than 4% of added fluorescent albumin. Transmigration was monitored at 20 hours post-infection either by direct dark-field counting or using a flow cytometer; both approaches gave closely corresponding values. At that time, less than 10% of a non-infectious (NI), non-adherent control strain (GCB705) transmigrated; in contrast, almost 55% of wild-type spirochetes (GCB726) did, for both types of primary cells (**Fig. 6B**). This highly efficient endothelial transmigration system provided a valuable tool for additional experiments as described below.

**Fig. 6.**
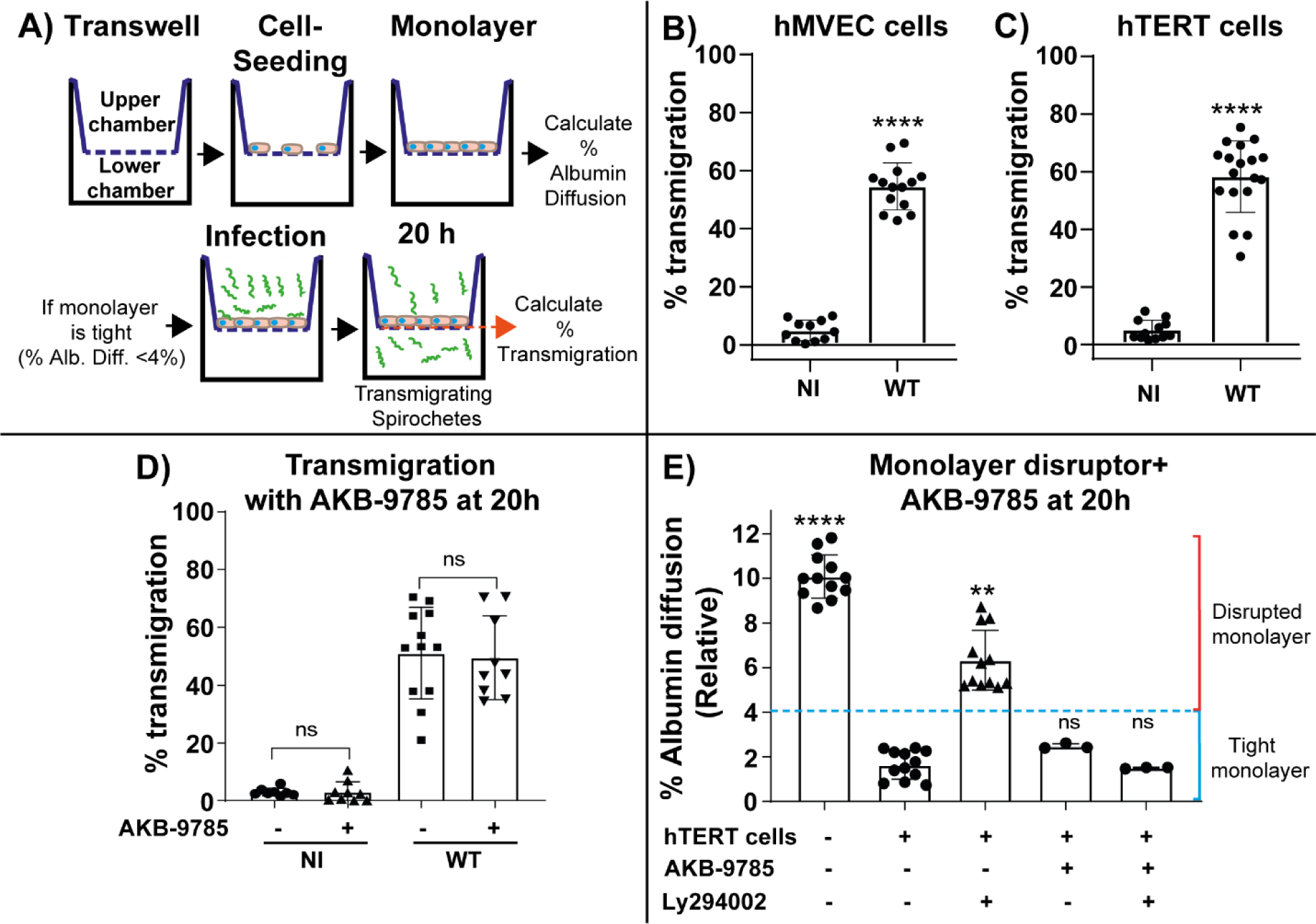
Transendothelial migration of *B. burgdorferi* in Transwell chambers. **A)** To study *B. burgdorferi* transmigration, Transwell chambers were seeded with hMVEC-d or hTERT cells until a tight monolayer was formed (<4% albumin diffusion). The upper chamber was infected with 3x10^5^ spirochetes, and the percentage of total transmigrated spirochetes (lower chamber) was determined after 20 h of infection by counting spirochetes in both the upper and lower chamber using a Petroff-Hauser chamber and dark field microscopy or by flow cytometry as indicated in each case. **B)** and **C)** The graphs show the percentage (mean ± SD) of *B. burgdorferi* that had transmigrated through human microvascular endothelial cells as determined by flow cytometry. The data represent the mean +/- SD of three independent experiments performed in quadruplicate. The p-value (p < 0.0001) was determined using the Mann-Whitney test. NI denotes the non-infectious strain GCB705, and WT indicates GCB726. **D)** Evaluation of *B. burgdorferi* transendothelial migration in hTERT treated with AKB-9785. The complete chamber was treated with 5 µM AKB9785b for the duration of the assay. The upper chamber was infected with 3x10^5^ spirochetes, and the percentage of total transmigrated spirochete was assessed by counting in a Petroff-Hausser chamber after 20 h of infection. The data represent the mean +/- SD of three independent experiments performed in quadruplicate; non-parametric ANOVA was performed to compare the control cell with the treatments using Kruskal-Wallis post-test (ns = not significant). **E)** Demonstration of the effectiveness of AKB-9785 to lock intercellular junctions in hTERT cells. Ly 294002 at 40 µM was used to disrupt the monolayer, and AKB-9785 was used to lock intercellular junctions and preserve monolayer integrity, which was assessed using an albumin diffusion assay: 10 ug of 555-Alb was added to the upper chamber at 16 h and the reading was carried out at the final point (20 h). The graph represents the mean +/- SD of three experiments performed in triplicate and measured in duplicate. Non-parametric ANOVA was performed to compare the control cell with the treatments using Kruskal-Wallis post-test; (p < 0.05) was considered significant.

Our results suggested that *B. burgdorferi* might transmigrate directly through cells, in agreement with our recent *in* vivo experiments indicating a transcellular pathway for vascular escape of *B. burgdorferi* in the mouse knee joint vasculature (Tan et al., 2021). Endothelial cells control permeability by modulating the phosphorylation level of adhesion molecules and/or their associated components (Bazzoni and Dejana, 2004; Daniel and Reynolds, 1997; Lampugnani et al., 1997). Many phosphatases and kinases are involved in this process. In general, a high level of phosphorylation promotes junction disassembly and opening of a paracellular pathway (Bogatcheva et al., 2002; Dominguez et al., 2007; Shi et al., 1998; Sui et al., 2005; Volberg et al., 1992; Young et al., 2003). Endothelial protein tyrosine phosphatase (VE-PTP) is exclusively expressed in endothelial cells (Baumer et al., 2006; Dominguez et al., 2007). Therefore, to distinguish between a paracellular and a transcellular pathway for spirochete transmigration in the Transwell chamber assay, we used AKB-9785 (Braun et al., 2019; Gurnik et al., 2016; Oehlers et al., 2016), an inhibitor of VE-PTP to answer this question. This inhibitor locks endothelial cell junctions. We treated hTERT cells with AKB-9785 (5μM) and infected them concomitantly with *B. burgdorferi* for 20h. Interestingly, the percentage of total transmigrated wild-type spirochetes through a human hTERT monolayer did not significantly decrease when the endothelial junctions were locked with AKB-9785 (**Fig. 6D**). We also performed an albumin diffusion control to demonstrate that AKB-9785 was effectively locking the junctions in the human hTERT cell monolayer. To do this, we used LY294002 (a PI3K and related proteins inhibitor, (Gharbi et al., 2007; Vlahos et al., 1994), which disrupted the monolayer, and AKB-9785 to successfully revert this effect and preserve the monolayer integrity, demonstrating the inhibitor’s effectiveness in our model (**Fig. 6E**).

Since the transmigration of *B. burgdorferi* was not affected by the locking of the cell-cell junctions, our results suggest a transcellular mechanism for transendothelial migration *in vitro* in human telomerase immortalized dermal cells, in agreement with our previous results in living mouse knee joint vasculature (Tan et al., 2021).

### Analysis of the roles of *B. burgdorferi* adhesins in the *in vitro* transendothelial migration system

It is known that *in vivo*, the first step in crossing the vasculature is adhesion. We have previously assessed the role of a number of adhesins in transmigration in living mice, including BBK32 (Moriarty et al., 2012; Norman et al., 2008), VlsE (Tan et al., 2022), P66 (Kumar et al., 2015), OspC (Lin et al., 2020) and DbpA/B (Tan et al., 2023); P66, DbpA and OspC were found to be required. Using endothelial cells and the Transwell chamber assay (**Fig. 6**) we studied several adhesin mutants (BBK32, VlsE, DbpA/B, OspC, and P66) and their effect on the transmigration process. As shown in (**Fig. S2**), surprisingly, none of the adhesin mutants alone adversely affected *B. burgdorferi* transendothelial migration.

### Endocytosis analysis

Since *B. burgdorferi* crosses the monolayer even when junctions were locked, we questioned whether a facilitated pathway might be involved in spirochete uptake. Previous findings in phagocytic cells showed that the actin cytoskeleton was involved in *B. burgdorferi* internalization, for example the inhibition of the microfilament reorganization significantly reduced internalization but not adhesion of the spirochete (Naj et al., 2013; Naj and Linder, 2017; Wu et al., 2011). However, the uptake mechanism in non-phagocytic cells has not been described. We performed an exploratory analysis by TEM (**Fig. 7A**) and found evidence of spirochetes adhered to the cell membrane (**Fig. 7A** left panel), cell membrane invagination in the contact point with the spirochetes (**Fig. 7A**, 2 middle panels) and spirochete penetration (**Fig. 7A**, 2 right panels). These findings were encouraging but did not point toward the pathway that might be involved in pathogen uptake. Therefore, to advance our understanding of this process, we selected inhibitors to block the most common endocytosis pathways (**Fig. 7B**). First, we inhibited dynamin formation with Dynasore (**Fig. 7B** left panel), which inhibits GTPase-dynamin-related endocytosis such as clathrin-dependent, caveolin-dependent (Henley et al., 1998), FEME (Fast endophilin-mediated endocytosis) (Casamento and Boucrot, 2020) and Rho/IL-2R (Basquin et al., 2013; Preta et al., 2015). Next, we used Filipin III, because it binds specifically to un-esterified cholesterol present in plasma membrane lipid rafts blocking flotillin-dependent and caveolin-dependent pathways. Finally, we used amiloride, which blocks the recruitment of Cdc42/Rac1 to the membrane by reducing the submembranous pH, and inhibits macropinocytosis (Koivusalo et al., 2010). Of the endocytic inhibitors tested, only amiloride diminished transmigration, with a decrease of 65% compared with the untreated control (**Fig. 7C**). Therefore, to ensure that this was not due to effects on spirochete growth, metabolic activity of the cells, or monolayer disruption, we tested the impact of amiloride on *B. burgdorferi* growth (**Fig. S3A**), in metabolic activity of the cells (**Fig. S3B**), and in monolayer disruption (**Fig. S3C**). Our results showed that amiloride does not modify spirochete growth nor the cell’s metabolic activity and does not disrupt the cell monolayer. Additionally, to confirm amiloride’s effect on macropinocytosis, we used dextran conjugated with tetramethylrhodamine. Dextran is one of the most used macropinosome markers because 70 kDa dextran is more selective for labelling of macropinosomes than 10 kDa dextran (Li et al., 2015). We quantified the particle area (macropinosomes) per total number of nuclei (**Fig. S3, D-E**). Macropinosome formation was reduced by 34% in 1 h 30 min with amiloride treatment. We hypothesize that this decrease is likely to be responsible for the 65% reduction in transmigration (**Fig. S3, D-E**).

**Fig. 7.**
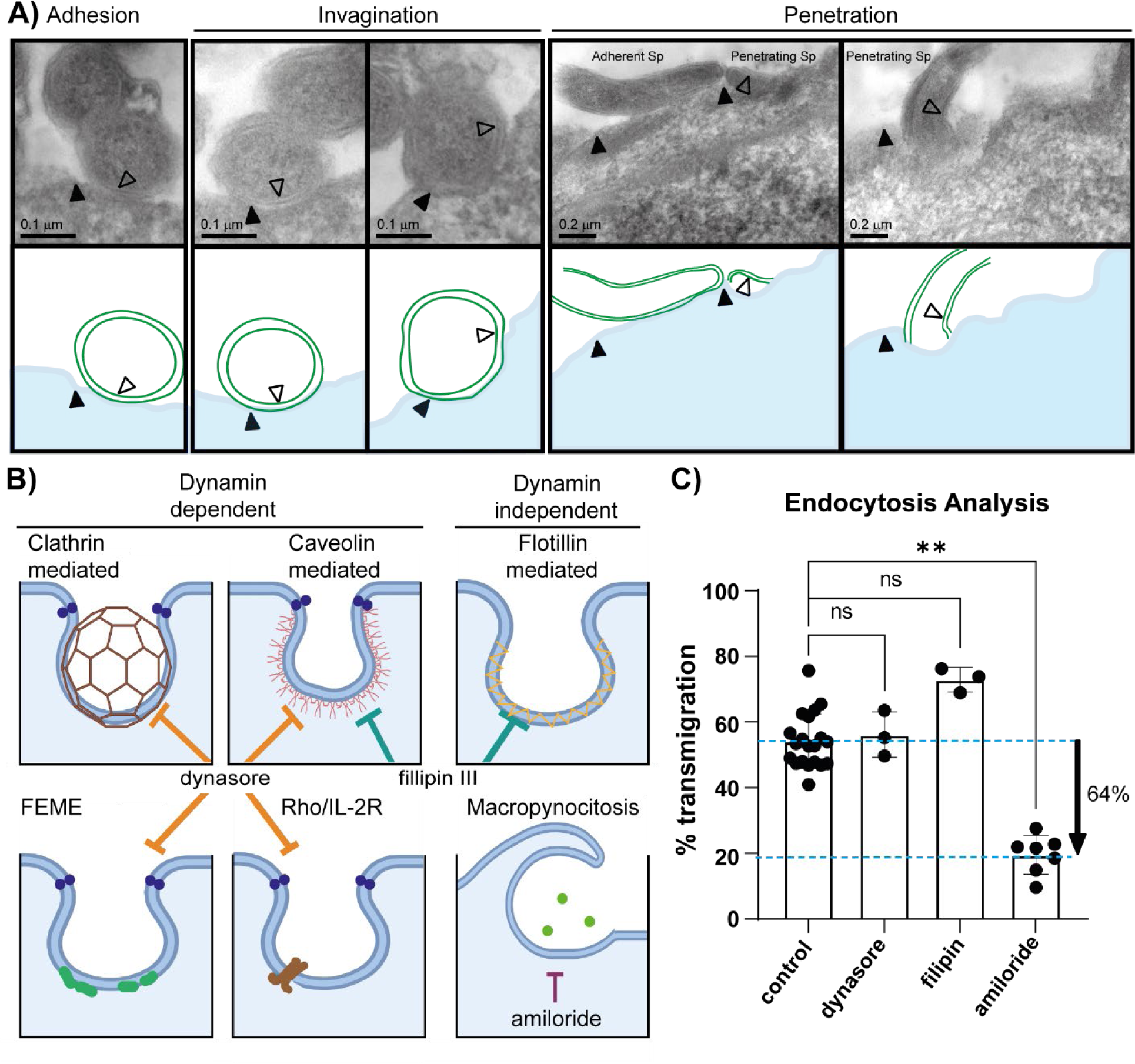
Transmigration analysis with endocytosis inhibitors. A) Transmission electron micrographs of *B. burgdorferi* in hMVEC cells (5 upper panels). The hMVEC cells were incubated with *B. burgdorferi* for 20 h in CoMe and then fixed with glutaraldehyde 2.5 %, embedded and sectioned for electron microscopy. The sections were observed using a Hitachi 7650 microscope, AMT16000 pixels camera. Spirochetes were captured in different stages of the penetration process: adherent to the cell surface (left), during the invagination process (2 central panels) as well as penetrating (2 right panels). Filled triangles indicate the cell membrane and empty triangles the double membrane of *B.burgdorferi*. The lower panels are schematics of the transmission electon micrographs. B) Schematic of some of the mechanisms of endocytosis and targets of the inhibitors used here. Amiloride is used to block macropinocytosis, dynasore rapidly and reversibly blocks dynamin-dependent endocytosis mediated by clathrin and caveolin and Filipin III blocks endocytosis mediated by caveolin and lipid rafts. C) Evaluation of B. burgdorferi transmigration with different endocytosis inhibitors using the Transwell assay. The upper chamber was infected with 3x10^5^ spirochetes and not treated (control) or treated with the indicated inhibitor: dynasore 80 μM, filipin 5 μM, and amiloride 6 μM. The data represent the mean +/- SD of three independent experiments performed in quadruplicate. Non-parametric ANOVA was performed to compare the untreated control with the various treatments using the Kruskal-Wallis post-test (A single * indicates *p* < 0.0332; ** indicates *p* < 0.0021; *** indicates *p* < 0.0002; **** indicates *p* < 0.0001).

### Internalization and transmigration analysis

The Rho family GTPases (Rho, Rac and Cdc42) are key proteins involved in several functions, including actin cytoskeletal organization, vesicle trafficking, and cell-to-cell and cell-to-extracellular matrix adhesions (Ridley, 2006). Therefore, we analyzed the internalization of *B. burgdorferi* (**Fig. 8, A-C**) and transmigration (**Fig. 8D**) by flow cytometry using amiloride to block the recruitment of Cdc42/Rac1 to the cell membrane. Amiloride treatment of endothelial cells resulted in an approximately 30% reduction in cell infection (**Fig. 8B**) and approximately 20% reduction in bacterial load or spirochete accumulation (**Fig 8C**).

**Fig. 8.**
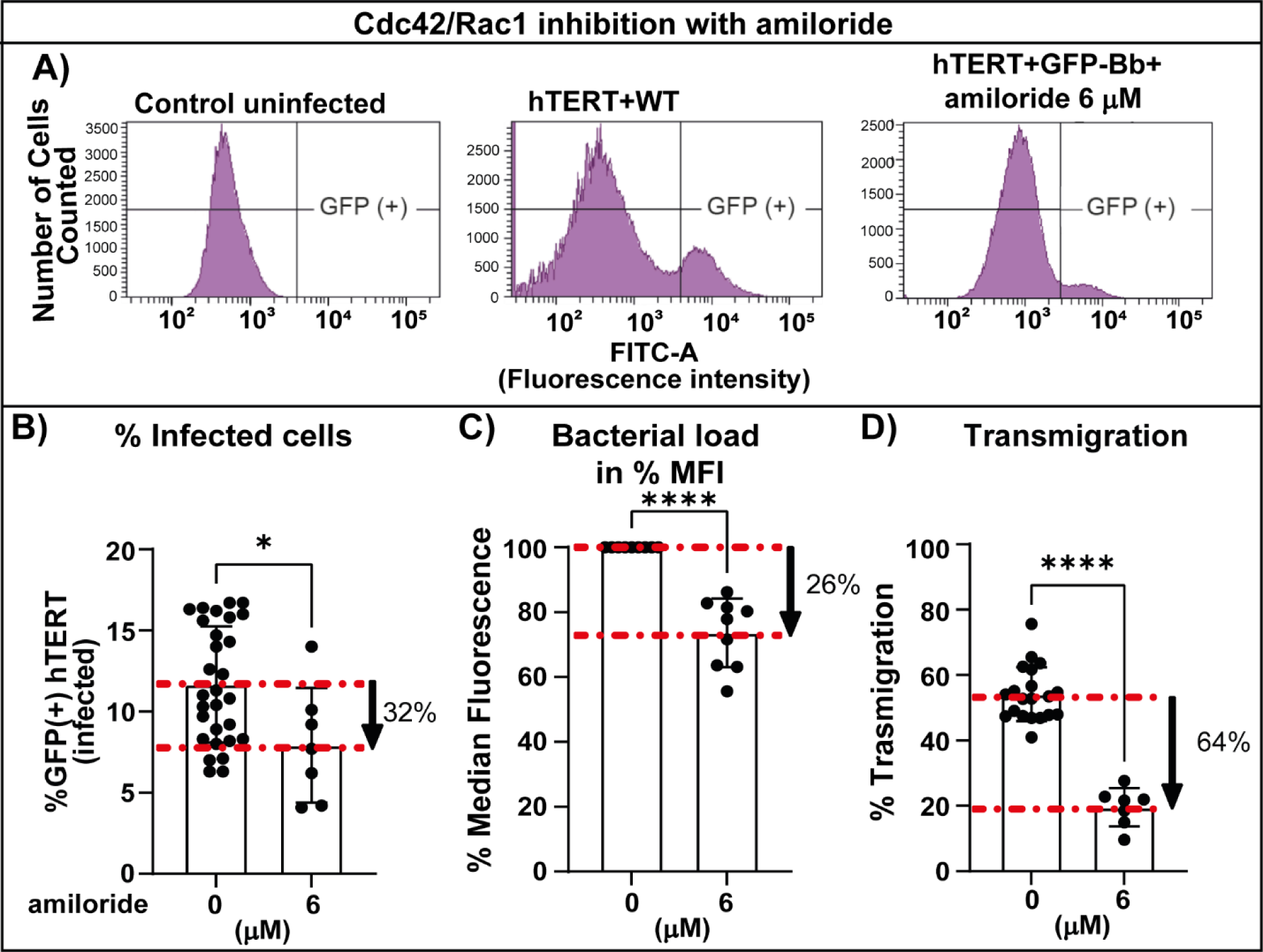
Analysis of the effect of Cdc42/Rac1 inhibitors on internalization, bacterial load and transendothelial migration. hTERT cells were grown in T-25 flasks and infected with *B. burgdorferi* GCB726 at a multiplicity of 7. After 20 h, the cells were treated with trypsin, washed with HBSS, fixed with PFA at 0.25% and run on the flow cytometer (1x10^5^ cells were analyzed). Statistical analysis was conducted in all cases using the Mann-Whitney test. A single * indicates *p* < 0.0332; ** indicates *p* < 0.0021; *** indicates *p* < 0.0002; **** indicates *p* < 0.0001. A) Representative plots of *B. burgdorferi* internalization into hTERT cells. Uninfected cells (left panel) were used to gate the GFP negative setting for the analysis. hTERT cells were infected with WT GFP-expressing *B. burgdorferi* alone or in the presence of 6 mm amiloride for 20 h, as indicated above the plots. B) The graph shows the percentage of cells infected with GFP-expressing *B. burgdorferi* in the absence or presence of 6 mM amiloride for 20 h. C) The MFI (median fluorescence intensity) was also quantified for each analyzed sample. The level of fluorescence gives an indication of bacterial load per cell. The experiments were carried out in triplicate, analyzing 2-3 samples per experiment. The data represent the mean +/- SD. D) Evaluation of *B. burgdorferi* transmigration with amiloride using the Transwell assay. The upper chamber was infected with 3x10^5^ spirochetes and treated with 6 μM amiloride. The data represent the mean +/- SD of three independent experiments performed in quadruplicate.

Next, to evaluate the effect of Cdc42 on *B. burgdorferi* internalization, we used ML141, an allosteric, reversible and non-competitive inhibitor that selectively blocks Cdc42 over other members of the Rho family at doses lower than 100 µM (Hong et al., 2013). Similarly to amiloride, blocking Cdc42 reduced the percentage of infected cells by 34% (**Fig. 9, A-B**) (compared to 32% found in amiloride treated cells) and bacterial load by 24% (**Fig. 9C**) (compared to 26% in amiloride treatment); suggesting that Cdc42 may be necessary for *B.burgdorferi* internalization in microvascular endothelial cells.

**Figure 9:**
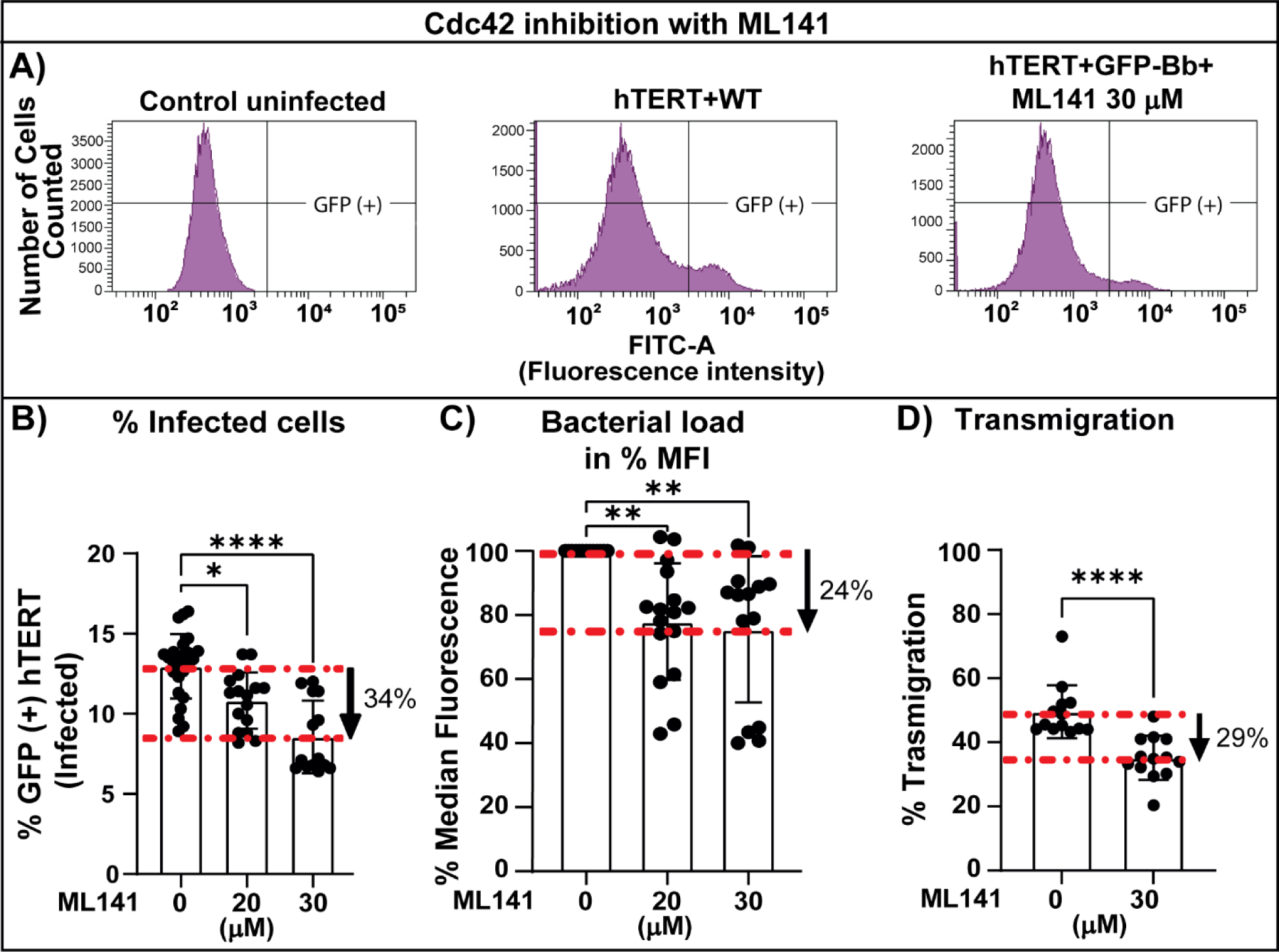
Analysis of the effect of Cdc42 inhibitor ML141 on internalization, bacterial load and transendothelial migration. hTERT cells were grown in T-25 flasks and infected with *B. burgdorferi* GCB726 at a multiplicity of 7. After 20 h, the cells were treated with trypsin, washed with HBSS, fixed with PFA at 0.25% and run on the flow cytometer (1x10^5^ cells were analyzed). Statistical analysis was conducted in all cases using the Mann-Whitney test. A single * indicates *p* < 0.0332; ** indicates *p* < 0.0021; *** indicates *p* < 0.0002; **** indicates *p* < 0.0001. A) Representative plots of *B. burgdorferi* internalization into hTERT cells. Uninfected cells (left panel) were used to gate the GFP negative setting for the analysis. hTERT cells were infected with WT GFP expressing *B. burgdorferi* alone, or in the presence of 30 µm ML141 for 20 h, as indicated above the plots. B) The graph shows the percentage of cells infected with GFP-expressing *B. burgdorferi* (from the initial 1x10^5^ cells analyzed) in the absence or presence of 20-30 µM ML141 for 20h. C) The MFI (median fluorescence intensity) was quantified from the positive (infected population) for each analyzed sample, thusthe level of fluorescence gives an indication of bacterial load per cell. The experiments were carried out in triplicate, analyzing 2-3 samples per experiment. The data represent the mean +/- SD. D) Evaluation of *B. burgdorferi* transmigration with ML141 using the Transwell assay. The upper chamber was infected with 3x10^5^ spirochetes and treated with 30 μM ML141, the dots represent the percentage of transmigrated spirochetes. The data represent the mean +/- SD of three independent experiments performed in quadruplicate.

In contrast, when analyzing transmigration (**Fig. 9D**), blocking Cdc 42 had a less marked effect, 29 % compared to 64 % found with amiloride, suggesting that Cdc42 is only partially responsible for the reduction in transmigration, pointing toward a role for Rac1 in the process. We confirmed that ML141 effects were not due to changes on *B. burgdorferi* growth (**Fig.S4A**), nor the metabolic activity of the cells (**Fig. S4B**), or a monolayer disruption (**Fig. S4C**).

We then evaluated the role of Rac1 using the internalization and transmigration assay (**Fig. 10, A-C**). NSC 23766 is a selective inhibitor of the interaction of Rac1 with guanine nucleotide exchange factors (GEFs). The inhibitor prevents Rac1 activation by Rac-specific TrioN and Tiam1 without affecting Cdc42 or RhoA activation (Gao et al., 2004). Blocking Rac1 alone had no effect on the percentage of cells infected (**Fig. 10B**), but significantly increased the bacterial accumulation by 97% (**Fig. 10C**), while reducing transmigration by 84.5% (**Fig. 10D**). In sum, this suggests that Rac1 is not involved in spirochete uptake, but is necessary for cellular egress. NSC 23766 treatment did not affect spirochete growth (**Fig.S5A**), metabolic activity of the cells (**Fig. S5B**), or the monolayer integrity (**Fig. S5C**).

**Figure 10:**
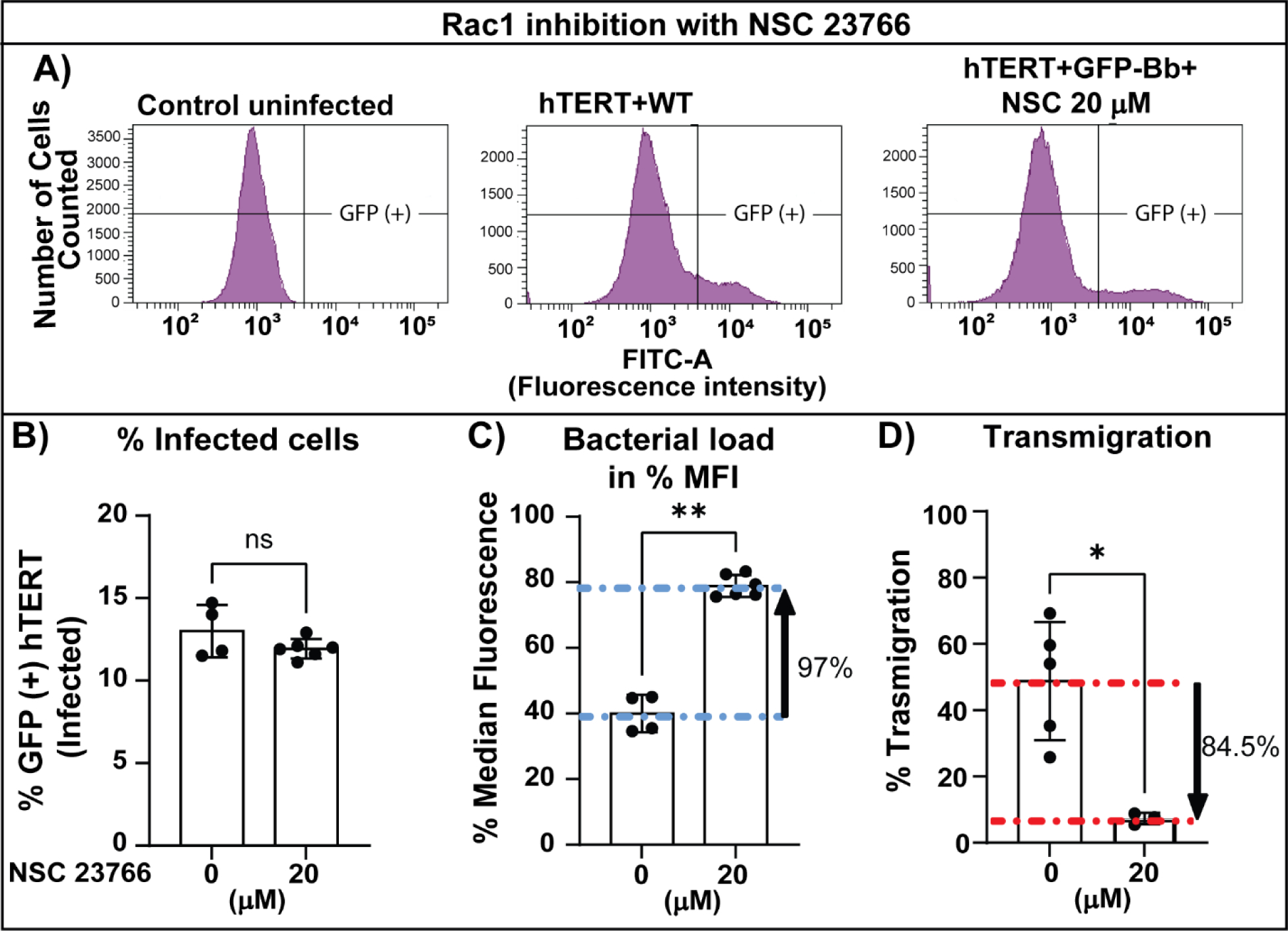
Analysis of the effect of Rac1 inhibitor NSC 23766 on internalization, bacterial load and transendothelial migration. hTERT cells were grown in T-25 flasks and infected with *B. burgdorferi* GCB726 at a multiplicity of 7. After 20 h, the cells were treated with trypsin, washed with HBSS, fixed with PFA at 0.25% and run on the flow cytometer (1x10^5^ cells were analyzed). Statistical analysis was conducted in all cases using the Mann-Whitney test. A single * indicates *p* < 0.0332; ** indicates *p* < 0.0021; *** indicates *p* < 0.0002; **** indicates *p* < 0.0001. A) Representative plots of *B. burgdorferi* internalization into hTERT cells. Uninfected cells (left panel) were used to gate the GFP negative setting for the analysis. hTERT cells were infected with WT GFP-expressing *B. burgdorferi* alone or in the presence of 20 µm NSC 23766 for 20 h as indicated above the plots. B) The graph shows the percentage of cells infected with GFP-expressing *B. burgdorferi* in the absence or presence of 20 µM NSC 23766 for 20h. C) The MFI (median fluorescence intensity) was also quantified for each analyzed sample. The level of fluorescence gives an indication of bacterial load per cell. The experiments were carried out in triplicate, analyzing 2-3 samples per experiment. The data represent the mean +/- SD. D) Evaluation of *B. burgdorferi* transmigration with NSC 23766 using the Transwell assay. The upper chamber was infected with 3x10^5^ spirochetes and treated with the indicated inhibitor. The data represent the mean +/- SD of three independent experiments performed in quadruplicate.

We also tested inhibitors that block various effectors, including PI3K, MEK1/3, Arp2/3 complex and p38, and Cdc42 and Rac 1/3 **(Table S3)**. However, in most cases, the inhibitors also affected the cells’ metabolic activity or permeability of the monolayer. From the tested inhibitors, only amiloride (Cdc42 and Rac1 membrane recruitment inhibitor) and ML141 (Cdc42 inhibitor) diminished internalization, bacteria load and transmigration without affecting *B. burgdorferi* growth or the metabolic activity of the cells. NSC 23766 (Rac1 inhibitor) did not affect cell infection but increased bacterial load and decreased transmigration. Other inhibitors (**Table S3**) were not informative.

## DISCUSSION

### Efficient transendothelial migration in human microvascular endothelial cells *in vitro*

Studying the mechanistic features of *B. burgdorferi* transmigration *in vivo* is difficult because it is a relatively rare event that is influenced by a variety of factors including blood flow, *B. burgdorferi* adhesins and gene expression, the immune response, and endothelial activation (Kumar et al., 2015; Lin et al., 2020; Moriarty et al., 2008; Moriarty et al., 2012; Norman et al., 2008; Tan et al., 2023; Tan et al., 2022; Tan et al., 2021). Therefore, we pursued *in vitro* studies where more controlled, detailed and variable analyses are possible (Ebady et al., 2016; Niddam et al., 2017). We created a suitable co-medium that promotes the survival of both *B. burgdorferi* and primary human dermal microvascular endothelial cells for 48 hours, allowing healthy maintenance of both cells and spirochetes for an adequate time to enable highly efficient transmigration in Transwell chambers, something not previously reported (**Table S1**). We routinely observed about 55% *B. burgdorferi* transendothelial migration at 20 hours post-infection in primary human dermal microvascular endothelial cells (hMVEC-d) and neonatal immortalized dermal microvascular endothelial cells (hTERT). Previous findings have shown that the spirochetes travel from the apical to the basolateral surface, indicating that the spirochetes can sense the polarity of the endothelial surface in the monolayer (Hirschberg et al., 2005; Zovein et al., 2010). Therefore, because of the system’s efficiency, facile enumeration can be performed by dark field microscopy or flow cytometry.

In contrast to results from transmigration studies in the mouse knee-joint using intravital imaging, we did not find a need for the *B. burgdorferi* adhesins P66 (Kumar et al., 2015), OspC (Lin et al., 2020) or DbpA (Tan et al., 2023) *in vitro* (**Fig. S2**). This somewhat surprising finding may result from substantial differences between our cell culture system and the mouse infection model. In the former, there is an obvious lack of blood, shear forces resulting from blood flow, an active immune system, secreted immune mediators, pericytes, and a basement membrane. Moreover, the *in vitro* system utilizes cells that are maintained in a growing state due to a cell medium rich in serum and growth factors (hFGF, hEFG, VEGF, IGF), which may enhance membrane trafficking/endocytosis (Barbieri et al., 2000; Brunk et al., 1976; Eichmann and Simons, 2012; Simons, 2012; Wang et al., 2019). Finally, important differences no doubt exist between the human dermal cells used here and endothelial cells found in the mouse knee joint periphery. Yet, an important point of congruence between our *in vitro* system and intravital imaging in the mouse knee joint is that the non-infectious, non-adherent strain GCB705, which lacks a variety of adhesins, is unable to cross the endothelium *in vitro* (**Fig. 6**) or *in vivo* (Norman et al., 2008).

Moreover, a transcellular rather than paracellular route for transmigration was observed both *in vitro* (**Fig. 3, 5**) and *in vivo* (Tan et al., 2021), suggesting that the underlying mechanism for transmigration may be similar in both cases. We believe that the pursuit of further *in vitro* studies may reveal significant new mechanistic findings that are inaccessible in the more cumbersome whole animal experimental system and that further studies will clarify the differences in the requirement for several *B. burgdorferi* adhesins exist *in vivo* versus *in vitro*.

### Spirochete penetration and transmigration studies support a transcellular pathway for transmigration

Until now, the characterization of spirochete penetration and internalization has not been achievable *in vivo* or *in vitro.* Studies thus far have been limited to the observation of internalized spirochetes in a variety of cells, for example, neutrophils, macrophages, neuroglia, fibroblast and macroendothelial cells (Livengood and Gilmore Jr, 2006; Petnicki-Ocwieja and Kern, 2014; Williams et al., 2018; Wu et al., 2011). Here we report internalization of spirochetes in primary human dermal microvascular endothelial cells to levels of 24 percent at 16 hours post-infection (**Fig. 3**).

Furthermore, we visualized penetrating spirochetes and found that internalization was occurring at a wide variety of locations on the cell surface rather than specifically near the cellular boundaries (**Fig. 5**). The widespread cellular localization of penetrating *B. burgdorferi* is expected if these spirochetes go on to transmigrate using a transcellular pathway.

Early *in vitro* studies on the pathways used by *B. burgdorferi* for extravasation reported conflicting results, favoring both paracellular and transcellular transmigration (Comstock and Thomas, 1989, 1991; Szczepanski et al., 1990). More recently, *in vivo* studies using intravital imaging coupled with AKB-9785, an inhibitor of the vascular endothelial protein tyrosine phosphatase (VE-PTP) that locks endothelial junctions (Braun et al., 2019; Gurnik et al., 2016; Oehlers et al., 2016), supported a transcellular pathway for extravasation into tissue surrounding the mouse knee joint (Tan et al., 2021). Here, we also used AKB-9785 with our *in vitro* transmigration system to investigate the transmigration pathway (**Fig. 6D, E**). AKB-9785 was shown to indeed lock endothelial junctions in the monolayer, and as we previously observed *in vivo* (Tan et al., 2021), it did not decrease *B. burgdorferi* transmigration levels in our Transwell chambers. Our results further support a transcellular pathway involved in *B. burgdorferi* transmigration when traversing human dermal microvascular endothelial cells, in agreement with our previous observations *in vivo* in mouse knee joint (Tan et al., 2021).

### A role for Cdc42 and Rac1 on *B. burgdorferi* internalization and transmigration: internalized spirochetes are precursors in the transendothelial migration pathway

Since *B. burgdorferi* was crossing the monolayer even when junctions were locked, we wondered whether a facilitated cellular pathway might be involved in spirochete uptake. Previous findings in phagocytic cells showed that the actin cytoskeleton was involved in *B. burgdorferi* internalization (Naj et al., 2013; Naj and Linder, 2017; Wu et al., 2011). Studies using other spirochetes have revealed differences in endothelial migration mechanisms. *T. pallidum* may use a cholesterol-dependent mechanism for endocytosis and also disrupts VE-cadherin intercellular junctions (Lithgow et al., 2021). Similarly, *Leptospira interrogans* also appears to promote dual pathways, but involving metallopeptidases to hydrolyze junction proteins (Ge et al., 2020) or microfilament-dependent endocytosis (Zhao et al., 2022). In contrast, *Neisseria meningitidis* invades endothelial cells by triggering the formation of membrane protrusions, leading to bacterial uptake using the Rac1 signalling pathway (Lambotin et al., 2005). To help distinguish between the alternative pathways shown in (**Fig. 7B**), we first inhibited dynamin formation with dynasore, lipid raft formation with filipin III, and recruitment of Cdc42/Rac1 to the membrane with amiloride (Koivusalo et al., 2010). In this first approach, only amiloride diminished transmigration, with a decrease of 65% compared with the untreated control without disrupting *B. burgdorferi* growth, the cell’s metabolic activity or the cell monolayer.

The amiloride family inhibits the activity of Na+/H+ exchangers, leading to acidification of the sub-membranous cytosol with a subsequent failure in recruitment of the small GTPases Rac1 and Cdc42 to the plasma membrane (Koivusalo et al., 2010). A limitation of using amiloride-derived inhibitors is that they are in fact inhibitors of Na+/H+, Na+, and Na+/Ca2+ exchangers (Orlowski and Grinstein, 1997); therefore, we used more specific inhibitors toward Cdc42 and Rac1 with varying chemical structures (**Fig. S6**) to clarify their roles in this process. ML141, at the doses used in our study, is a specific inhibitor of Cdc42 with no activity toward Ras and Rac1 (Hong et al., 2013); the results with ML141 showed a reduction in both internalization and transmigration without affecting spirochete growth, cell metabolic activity of the cells, or the monolayer permeability (**Fig. 9**). Moreover, using the specific NSC 23766 Rac1 inhibitor, which has no activity towards Cdc42 or RhoA (Gao et al., 2004), we observed a dramatic increase in bacterial load coupled with a dramatic decrease in transmigration (**Fig. 10**). These compelling results strongly suggest that Rac1 is a crucial component of the cellular egress step of transendothelial migration and that spirochete internalization is a precursor to cellular exit. The chances that the three drugs used here, with very different chemical structures (**Fig. S6**) all perturb spirochete transendothelial migration through off target effects are minute. Our data from experiments performed in the absence of inhibitors (**Fig. 8-10**) also suggest that cellular egress is the rate-limiting step in the transendothelial migration pathway as internalized *B. burgdorferi* accumulate at 20 hours in the absence of drug treatment.

Different bacterial pathogens, including *Legionella pneumophila*, *Burkholderia cenocepacia, Salmonella typhimurium* (Rosales-Reyes et al., 2018; Watarai et al., 2001), *Mycobacterium tuberculosis* and *Mycobacterium smegmatis* (García-Pérez et al., 2012; García-Pérez et al., 2008) invade cells by modulating Cdc42 and Rac1 via virulence factors or by using an uncommon set of receptors (Popoff and Geny, 2009; Rottner et al., 2005). No orthologues of these factors exist in *B. burgdorferi*, which suggests that the use of Cdc42 and Rac1 by the Lyme disease spirochete occurs through a previously undescribed mechanism. A remaining question is to distinguish between the actual use of the macropinocytosis pathway versus a simple requirement for the Cdc42 and Rac1 GTPases. Future studies in the molecular mechanisms involved in transendothelial migration by *B. burgdorferi* will benefit from the co-culture conditions described here.

## Acknowledgements

Research reported in this publication was supported by grants PJT-153336 and PJT-180584 from the Canadian Institutes of Health Research and by the National Institute of Allergy and Infectious Diseases of the National Institutes of Health under award numbers R01AI18799 and R01AI169724. Confocal microscopy imaging was performed in the Live Cell Imaging Laboratory, and flow cytometry was performed in the Nicole Perkins Microbial Communities Core facility, both at the Snyder Institute for Chronic Diseases at the University of Calgary. TEM was conducted in the Microscopy and Imaging Facility at the Cumming School of Medicine, University of Calgary. We are grateful to Dr. Kevin Peters at Aerpio Therapeutics for generously providing AKB-9785. We thank Karen Poon, Priyanka G. Mukherjee, Lucy Swift and Genevieve Chaconas for technical assistance.

## Author Contributions

Conceptualization: DA-O, CK, RD, and GC were responsible for the conceptual research idea and study design. Formal Analysis, investigation, methodology, project administration and validation: DA-O and CK. Visualization, DA-O and CK. Data curation/software design: TV wrote the script to analyze microscopy data. Funding acquisition, Resources: RD and GC. Supervision: MH, RD and GC. Writing – original draft preparation DA-O, RD and GC. Writing – review and editing: All authors have reviewed, edited and agreed to the published version of the manuscript.

## Notes

### Competing Interest Statement

The authors have declared no competing interest.

## References

Barbieri, M.A., Roberts, R.L., Gumusboga, A., Highfield, H., Alvarez-Dominguez, C., Wells, A., and Stahl, P.D. (2000). Epidermal growth factor and membrane trafficking: EGF receptor activation of endocytosis requires Rab5a. The Journal of cell biology 151, 539–550.

Barbour, A.G. (1984). Isolation and cultivation of Lyme disease spirochetes. Yale J Biol Med 57, 521–525.

Barbour, A.G. (1986). Cultivation of Borrelia: a historical overview. Zentralbl Bakteriol Mikrobiol Hyg [A] 263, 11-14.

Basquin, C., Malardé, V., Mellor, P., Anderson, D.H., Meas-Yedid, V., Olivo-Marin, J.-C., Dautry-Varsat, A., and Sauvonnet, N. (2013). The signalling factor PI3K is a specific regulator of the clathrin-independent dynamin-dependent endocytosis of IL-2 receptors. Journal of cell science 126, 1099–1108.

Baumer, S., Keller, L., Holtmann, A., Funke, R., August, B., Gamp, A., Wolburg, H., Wolburg-Buchholz, K., Deutsch, U., and Vestweber, D. (2006). Vascular endothelial cell–specific phosphotyrosine phosphatase (VE-PTP) activity is required for blood vessel development. Blood 107, 4754–4762.

Bazzoni, G., and Dejana, E. (2004). Endothelial cell-to-cell junctions: molecular organization and role in vascular homeostasis. Physiological reviews 84, 869–901.

Bednarek, R. (2022). In vitro methods for measuring the permeability of cell monolayers. Methods and protocols 5, 17.

Bogatcheva, N.V., Garcia, J.G., and Verin, A.D. (2002). Role of tyrosine kinase signaling in endothelial cell barrier regulation. Vascular pharmacology 39, 201–212.

Bono, J.L., Elias, A.F., Kupko, J.J., III, Stevenson, B., Tilly, K., and Rosa, P. (2000). Efficient targeted mutagenesis in *Borrelia burgdorferi*. J Bacteriol 182, 2445–2452.

Braun, L.J., Zinnhardt, M., Vockel, M., Drexler, H.C., Peters, K., and Vestweber, D. (2019). VE-PTP inhibition stabilizes endothelial junctions by activating FGD5. EMBO Rep 20, e47046.

Brunk, U., Schellens, J., and Westermark, B. (1976). Influence of epidermal growth factor (EGF) on ruffling activity, pinocytosis and proliferation of cultivated human glia cells. Experimental cell research 103, 295–302.

Casamento, A., and Boucrot, E. (2020). Molecular mechanism of fast endophilin-mediated endocytosis. Biochemical Journal 477, 2327–2345.

Coburn, J., Garcia, B., Hu, L.T., Jewett, M.W., Kraiczy, P., Norris, S.J., and Skare, J. (2021). Lyme Disease Pathogenesis. Curr Issues Mol Biol 42, 473–518.

Comstock, L.E., and Thomas, D.D. (1989). Penetration of endothelial cell monolayers by *Borrelia burgdorferi*. Infect Immun 57, 1626–1628.

Comstock, L.E., and Thomas, D.D. (1991). Characterization of *Borrelia burgdorferi* invasion of cultured endothelial cells. Microb Pathog 10, 137–148.

Daniel, J.M., and Reynolds, A.B. (1997). Tyrosine phosphorylation and cadherin/catenin function. Bioessays 19, 883–891.

Dominguez, M.G., Hughes, V.C., Pan, L., Simmons, M., Daly, C., Anderson, K., Noguera-Troise, I., Murphy, A.J., Valenzuela, D.M., and Davis, S. (2007). Vascular endothelial tyrosine phosphatase (VE-PTP)-null mice undergo vasculogenesis but die embryonically because of defects in angiogenesis. Proceedings of the National Academy of Sciences 104, 3243–3248.

Ebady, R., Niddam, A.F., Boczula, A.E., Kim, Y.R., Gupta, N., Tang, T.T., Odisho, T., Zhi, H., Simmons, C.A., Skare, J.T., et al. (2016). Biomechanics of *Borrelia burgdorferi* Vascular Interactions. Cell Rep.

Eichmann, A., and Simons, M. (2012). VEGF signaling inside vascular endothelial cells and beyond. Current opinion in cell biology 24, 188–193.

Ferrara, N., Houck, K., Jakeman, L., and Leung, D.W. (1992). Molecular and biological properties of the vascular endothelial growth factor family of proteins. Endocrine reviews 13, 18–32.

Gao, Y., Dickerson, J.B., Guo, F., Zheng, J., and Zheng, Y. (2004). Rational design and characterization of a Rac GTPase-specific small molecule inhibitor. Proceedings of the National Academy of Sciences 101, 7618–7623.

García-Pérez, B.E., De la Cruz-López, J.J., Castañeda-Sánchez, J.I., Muñóz-Duarte, A.R., Hernández-Pérez, A.D., Villegas-Castrejón, H., García-Latorre, E., Caamal-Ley, A., and Luna-Herrera, J. (2012). Macropinocytosis is responsible for the uptake of pathogenic and non-pathogenic mycobacteria by B lymphocytes (Raji cells). BMC microbiology 12, 1–14.

García-Pérez, B.E., Hernández-González, J.C., García-Nieto, S., and Luna-Herrera, J. (2008). Internalization of a non-pathogenic mycobacteria by macropinocytosis in human alveolar epithelial A549 cells. Microbial pathogenesis 45, 1–6.

Ge, Y.M., Sun, A.H., Ojcius, D.M., Li, S.J., Hu, W.L., Lin, X., and Yan, J. (2020). M16-Type Metallopeptidases Are Involved in Virulence for Invasiveness and Diffusion of Leptospira interrogans and Transmission of Leptospirosis. J Infect Dis 222, 1008–1020.

Gharbi, S.I., Zvelebil, M.J., Shuttleworth, S.J., Hancox, T., Saghir, N., Timms, J.F., and Waterfield, M.D. (2007). Exploring the specificity of the PI3K family inhibitor LY294002. Biochemical Journal 404, 15–21.

Gurnik, S., Devraj, K., Macas, J., Yamaji, M., Starke, J., Scholz, A., Sommer, K., Di Tacchio, M., Vutukuri, R., Beck, H., et al. (2016). Angiopoietin-2-induced blood-brain barrier compromise and increased stroke size are rescued by VE-PTP-dependent restoration of Tie2 signaling. Acta Neuropathol 131, 753–773.

Henley, J.R., Krueger, E.W., Oswald, B.J., and McNiven, M.A. (1998). Dynamin-mediated internalization of caveolae. The Journal of cell biology 141, 85–99.

Hirschberg, R.M., Sachtleben, M., and Plendl, J. (2005). Electron microscopy of cultured angiogenic endothelial cells. Microscopy research and technique 67, 248–259.

Hong, L., Kenney, S.R., Phillips, G.K., Simpson, D., Schroeder, C.E., Nöth, J., Romero, E., Swanson, S., Waller, A., Strouse, J.J., et al. (2013). Characterization of a Cdc42 protein inhibitor and its use as a molecular probe. J Biol Chem 288, 8531–8543.

Kawabata, H., Norris, S.J., and Watanabe, H. (2004). BBE02 disruption mutants of *Borrelia burgdorferi* B31 have a highly transformable, infectious phenotype. Infect Immun 72, 7147–7154.

Koivusalo, M., Welch, C., Hayashi, H., Scott, C.C., Kim, M., Alexander, T., Touret, N., Hahn, K.M., and Grinstein, S. (2010). Amiloride inhibits macropinocytosis by lowering submembranous pH and preventing Rac1 and Cdc42 signaling. Journal of Cell Biology 188, 547–563.

Kumar, D., Ristow, L.C., Shi, M., Mukherjee, P., Caine, J.A., Lee, W.Y., Kubes, P., Coburn, J., and Chaconas, G. (2015). Intravital Imaging of Vascular Transmigration by the Lyme Spirochete: Requirement for the Integrin Binding Residues of the *B. burgdorferi* P66 Protein. PLoS Pathog 11, e1005333.

Lafrance, M.E., Pierce, J.V., Antonara, S., and Coburn, J. (2011). The *Borrelia burgdorferi* integrin ligand, P66, affects gene expression by human cells in culture. Infect Immun.

Lambotin, M., Hoffmann, I., Laran-Chich, M.P., Nassif, X., Couraud, P.O., and Bourdoulous, S. (2005). Invasion of endothelial cells by Neisseria meningitidis requires cortactin recruitment by a phosphoinositide-3-kinase/Rac1 signalling pathway triggered by the lipo-oligosaccharide. J Cell Sci 118, 3805–3816.

Lampugnani, M.G., Corada, M., Andriopoulou, P., Esser, S., Risau, W., and Dejana, E. (1997). Cell confluence regulates tyrosine phosphorylation of adherens junction components in endothelial cells. Journal of cell science 110, 2065–2077.

Leopold, B., Strutz, J., Weiß, E., Gindlhuber, J., Birner-Gruenberger, R., Hackl, H., Appel, H.M., Cvitic, S., and Hiden, U. (2019). Outgrowth, proliferation, viability, angiogenesis and phenotype of primary human endothelial cells in different purchasable endothelial culture media: Feed wisely. Histochemistry and Cell Biology 152, 377–390.

Li, L., Wan, T., Wan, M., Liu, B., Cheng, R., and Zhang, R. (2015). The effect of the size of fluorescent dextran on its endocytic pathway. Cell biology international 39, 531–539.

Lin, Y.P., Tan, X., Caine, J.A., Castellanos, M., Chaconas, G., Coburn, J., and Leong, J.M. (2020). Strain-specific joint invasion and colonization by Lyme disease spirochetes is promoted by outer surface protein C. PLoS Pathog 16, e1008516.

Lithgow, K.V., Tsao, E., Schovanek, E., Gomez, A., Swayne, L.A., and Cameron, C.E. (2021). *Treponema pallidum* Disrupts VE-Cadherin Intercellular Junctions and Traverses Endothelial Barriers Using a Cholesterol-Dependent Mechanism. Frontiers in microbiology 12, 691731.

Livengood, J.A., and Gilmore Jr, R.D. (2006). Invasion of human neuronal and glial cells by an infectious strain of Borrelia burgdorferi. Microbes and infection 8, 2832–2840.

Ma, Y., Sturrock, A., and Weis, J.J. (1991). Intracellular localization of *Borrelia burgdorferi* within human endothelial cells. Infect Immun 59, 671–678.

Moriarty, T.J., Norman, M.U., Colarusso, P., Bankhead, T., Kubes, P., and Chaconas, G. (2008). Real-time high resolution 3D imaging of the lyme disease spirochete adhering to and escaping from the vasculature of a living host. PLoS Pathog 4, e1000090.

Moriarty, T.J., Shi, M., Lin, Y.P., Ebady, R., Zhou, H., Odisho, T., Hardy, P.O., Salman-Dilgimen, A., Wu, J., Weening, E.H., et al. (2012). Vascular binding of a pathogen under shear force through mechanistically distinct sequential interactions with host macromolecules. Mol Microbiol 86, 1116–1131.

Naj, X., Hoffmann, A.-K., Himmel, M., and Linder, S. (2013). The formins FMNL1 and mDia1 regulate coiling phagocytosis of Borrelia burgdorferi by primary human macrophages. Infection and immunity 81, 1683–1695.

Naj, X., and Linder, S. (2017). Actin-dependent regulation of Borrelia burgdorferi phagocytosis by macrophages. The Actin Cytoskeleton and Bacterial Infection, 133-154.

Niddam, A.F., Ebady, R., Bansal, A., Koehler, A., Hinz, B., and Moriarty, T.J. (2017). Plasma fibronectin stabilizes *Borrelia burgdorferi*-endothelial interactions under vascular shear stress by a catch-bond mechanism. Proc Natl Acad Sci U S A 114, E3490–E3498.

Norman, M.U., Moriarty, T.J., Dresser, A.R., Millen, B., Kubes, P., and Chaconas, G. (2008). Molecular mechanisms involved in vascular interactions of the Lyme disease pathogen in a living host. PLoS Pathog 4, e1000169.

Nyarko, E., Grab, D., and Dumler, J. (2006). Anaplasma phagocytophilum-infected neutrophils enhance transmigration of Borrelia burgdorferi across the human blood brain barrier in vitro. International journal for parasitology 36, 601–605.

Oehlers, S.H., Cronan, M.R., Beerman, R.W., Johnson, M.G., Huang, J., Kontos, C.D., Stout, J.E., and Tobin, D.M. (2016). Infection-Induced Vascular Permeability Aids Mycobacterial Growth. J Infect Dis.

Orlowski, J., and Grinstein, S. (1997). Na+/H+ exchangers of mammalian cells. Journal of Biological Chemistry 272, 22373–22376.

Petnicki-Ocwieja, T., and Kern, A. (2014). Mechanisms of Borrelia burgdorferi internalization and intracellular innate immune signaling. Frontiers in cellular and infection microbiology 4, 175.

Popoff, M.R., and Geny, B. (2009). Multifaceted role of Rho, Rac, Cdc42 and Ras in intercellular junctions, lessons from toxins. Biochimica et Biophysica Acta (BBA)-Biomembranes 1788, 797-812.

Preta, G., Cronin, J.G., and Sheldon, I.M. (2015). Dynasore-not just a dynamin inhibitor. Cell Communication and Signaling 13, 1–7.

Ridley, A.J. (2006). Rho GTPases and actin dynamics in membrane protrusions and vesicle trafficking. Trends in cell biology 16, 522–529.

Ristow, L.C., Bonde, M., Lin, Y.P., Sato, H., Curtis, M., Wesley, E., Hahn, B.L., Fang, J., Wilcox, D.A., Leong, J.M., et al. (2015). Integrin binding by *Borrelia burgdorferi* P66 facilitates dissemination but is not required for infectivity. Cell Microbiol 17, 1021–1036.

Rosales-Reyes, R., Sánchez-Gómez, C., Ortiz-Navarrete, V., and Santos-Preciado, J.I. (2018). Burkholderia cenocepacia induces macropinocytosis to enter macrophages. BioMed research international 2018.

Rottner, K., Stradal, T.E., and Wehland, J. (2005). Bacteria-host-cell interactions at the plasma membrane: stories on actin cytoskeleton subversion. Developmental cell 9, 3–17.

Schindelin, J., Arganda-Carreras, I., Frise, E., Kaynig, V., Longair, M., Pietzsch, T., Preibisch, S., Rueden, C., Saalfeld, S., and Schmid, B. (2012). Fiji: an open-source platform for biological-image analysis. Nature methods 9, 676-682.

Shi, S., Verin, A.D., Schaphorst, K., Gilbert-McClain, L., Patterson, C., Irwin, R., Natarajan, V., and Garcia, J. (1998). Role of tyrosine phosphorylation in thrombin-induced endothelial cell contraction and barrier function. Endothelium 6, 153–171.

Simons, M. (2012). An inside view: VEGF receptor trafficking and signaling. Physiology 27, 213–222.

Stanek, G., and Strle, F. (2018). Lyme borreliosis-from tick bite to diagnosis and treatment. FEMS Microbiol Rev 42, 233-258.

Steere, A.C., Strle, F., Wormser, G.P., Hu, L.T., Branda, J.A., Hovius, J.W., Li, X., and Mead, P.S. (2016). Lyme borreliosis. Nat Rev Dis Primers 2, 16090.

Sui, X.F., Kiser, T.D., Hyun, S.W., Angelini, D.J., Del Vecchio, R.L., Young, B.A., Hasday, J.D., Romer, L.H., Passaniti, A., and Tonks, N.K. (2005). Receptor protein tyrosine phosphatase μ regulates the paracellular pathway in human lung microvascular endothelia. The American journal of pathology 166, 1247–1258.

Szczepanski, A., Furie, M., Benach, J., Lane, B., and Fleit, H. (1990). Interaction between *Borrelia burgdorferi* and endothelium *in vitro*. J Clin Invest 85, 1637–1647.

Tan, X., Castellanos, M., and Chaconas, G. (2023). Choreography of Lyme Disease Spirochete Adhesins To Promote Vascular Escape. Microbiol Spectr, e0125423.

Tan, X., Lin, Y.P., Pereira, M.J., Castellanos, M., Hahn, B.L., Anderson, P., Coburn, J., Leong, J.M., and Chaconas, G. (2022). VlsE, the nexus for antigenic variation of the Lyme disease spirochete, also mediates early bacterial attachment to the host microvasculature under shear force. PLoS Pathog 18, e1010511.

Tan, X., Petri, B., DeVinney, R., Jenne, C.N., and Chaconas, G. (2021). The Lyme disease spirochete can hijack the host immune system for extravasation from the microvasculature. Mol Microbiol 116, 498–515.

Thomas, D., and Comstock, L. (1989). Interaction of Lyme disease spirochetes with cultured eucaryotic cells. Infect Immun 57, 1324-1326.

Van Gundy, T.J., Ullmann, A.J., Brandt, K.S., and Gilmore, R.D. (2021). A transwell assay method to evaluate Borrelia burgdorferi sensu stricto migratory chemoattraction toward tick saliva proteins. Ticks and Tick-borne Diseases 12, 101782.

Vlahos, C.J., Matter, W.F., Hui, K.Y., and Brown, R.F. (1994). A specific inhibitor of phosphatidylinositol 3-kinase, 2-(4-morpholinyl)-8-phenyl-4H-1-benzopyran-4-one (LY294002). Journal of Biological Chemistry 269, 5241-5248.

Volberg, T., Zick, Y., Dror, R., Sabanay, I., Gilon, C., Levitzki, A., and Geiger, B. (1992). The effect of tyrosine-specific protein phosphorylation on the assembly of adherens-type junctions. The EMBO journal 11, 1733–1742.

von Lackum, K., and Stevenson, B. (2005). Carbohydrate utilization by the Lyme borreliosis spirochete, Borrelia burgdorferi. FEMS microbiology letters 243, 173–179.

Wang, H., He, J., Luo, Y., Mu, M., Guo, S., Shen, L., Qian, Z., Fang, Q., and Song, C. (2019). IGF-1 Promotes endocytosis of alveolar epithelial cells through PI3K signaling. Annals of Clinical & Laboratory Science 49, 3–8.

Watarai, M., Derre, I., Kirby, J., Growney, J.D., Dietrich, W.F., and Isberg, R.R. (2001). Legionella pneumophila is internalized by a macropinocytotic uptake pathway controlled by the Dot/Icm system and the mouse Lgn1 locus✪. The Journal of experimental medicine 194, 1081–1096.

Williams, S.K., Weiner, Z.P., and Gilmore, R.D. (2018). Human neuroglial cells internalize Borrelia burgdorferi by coiling phagocytosis mediated by Daam1. PLoS One 13, e0197413.

Wormser, G.P. (2006). Hematogenous dissemination in early Lyme disease. Wien Klin Wochenschr 118, 634–637.

Wormser, G.P., Brisson, D., Liveris, D., Hanincova, K., Sandigursky, S., Nowakowski, J., Nadelman, R.B., Ludin, S., and Schwartz, I. (2008). *Borrelia burgdorferi* genotype predicts the capacity for hematogenous dissemination during early Lyme disease. J Infect Dis 198, 1358–1364.

Wu, J., Weening, E.H., Faske, J.B., Hook, M., and Skare, J.T. (2011). Invasion of eukaryotic cells by *Borrelia burgdorferi* requires β1 integrins and Src kinase activity. Infect Immun 79, 1338–1348.

Wyss, C., and Ermert, P. (1996). Borrelia burgdorferi is an adenine and spermidine auxotroph. Microbial ecology in health and disease 9, 181–185.

Young, B.A., Sui, X., Kiser, T.D., Hyun, S.W., Wang, P., Sakarya, S., Angelini, D.J., Schaphorst, K.L., Hasday, J.D., and Cross, A.S. (2003). Protein tyrosine phosphatase activity regulates endothelial cell-cell interactions, the paracellular pathway, and capillary tube stability. American Journal of Physiology-Lung Cellular and Molecular Physiology 285, L63–L75.

Zhao, X., Guo, J., Jia, X., Yang, Y., Liu, L., Nie, W., and Fang, Z. (2022). Internalization of Leptospira interrogans via diverse endocytosis mechanisms in human macrophages and vascular endothelial cells. PLoS neglected tropical diseases 16, e0010778.

Zovein, A.C., Luque, A., Turlo, K.A., Hofmann, J.J., Yee, K.M., Becker, M.S., Fassler, R., Mellman, I., Lane, T.F., and Iruela-Arispe, M.L. (2010). β1 integrin establishes endothelial cell polarity and arteriolar lumen formation via a Par3-dependent mechanism. Developmental cell 18, 39–51.

