## Supplementary Material for "Transendothelial migration of the Lyme disease spirochete involves spirochete internalization as an intermediate step through a transcellular pathway that involves Cdc42 and Rac1"

**Published studies – HUVEC cells**

| Reference | Borrelia/Media | Co-culture (time) | Co-culture medium |
|---|---|---|---|
| (Thomas and Comstock, 1989) | BSK-II | 4h | M199-20% FBS |
| (Grab et al., 2009) | BSK-II 6% RS | 16h | M199-20%HiFBS |
| (Brissette et al., 2013) | BSK-II | 6 to 72h | DMEM |
| (Lafrance et al., 2011) | MKP Human serum | 3h, 6h, 24h | Cell medium without antibiotic |

**Published studies – Primary cells/cell lines**

**Table S1. Published media suitable for short-term co-culture of *B. burgdorferi* and HUVEC or primary cells.**

| Reference | *Borrelia* culture media | Co-culture time | Co-culture medium |
|---|---|---|---|
| (Wu et al., 2011) | BSK-II + 6% RS | 2 to 18h | DMEM/M199 +10%FBS |
| (Ma et al., 1991) | BSK-II | Up to 3d | No detailed |
| (Burns et al., 1997) | Serum free BSK | 4h/8h/12h/24h | M199–20% Heat inactivated FBS (HIFBS ) |
| (Burns and Furie, 1998) | BSK | 8h | M199 + 20% HIFBS |
| (Sellati et al., 1996) | BSK | 4h | M199 + 20% HIFBS |
| (Sellati et al., 1995) | Serum free BSK | 5h | M199+20% HIFBS+HEPES |
| (Coleman et al., 1995) | Serum free BSK | 1h | M199 |
| (Grab et al., 2005) | BSK II 10% RS | | M199 10% HiFBS |
| (Lazarus et al., 2008) | BSK-II 6% RS | 24h | RPMI or RPMI 10% FBS and 20% BSK-II medium (RPMI-B) |
| (Chung et al., 2013) | BSK-II 6% RS | Over night | RPMI-B |

**RS = rabbit serum**

**FBS = fetal bovine serum**

| GCB strain | Description | Missing plasmids | Source/ Reference |
|---|---|---|---|
| 705 | B31-A (Bono et al., 2000) +pTM61 *gent,gfp* | | (Moriarty et al., 2008) |
| 726 | B31 5A4 NP1 (*kan*) (Kawabata et al., 2004) + pTM61 *gent,gfp* | | (Moriarty et al., 2008) |
| 847 | B31-A3 (Elias et al., 2002) + pTM61 *gent,gfp,* clone 23 | | (Kumar et al., 2015) |
| 849 | B31-A3 (Elias et al., 2002) *Δp66:kan* (K04 C3-14) + pTM61-*strep,gfp*, clone 1 | | (Kumar et al., 2015) |
| 3003 | B31-A3 (Elias et al., 2002) K04 C3-14 + $p66^{D205A,D207A}$ *gent,* restored to chromosome clone 2-30 + pTM61-*strep,gfp*, clone 2-1 | | (Kumar et al., 2015) |
| 3007 | B31-A3 (Elias et al., 2002) *ΔospC::kan* +pTM61-*strep* | | (Lin et al., 2020) |
| 4032 | B31 5A4 (Purser and Norris, 2000) *ΔdbpA,B::gent,* pTM61-*kan, gfp* | cp9, lp21 | (Tan et al., 2023) |
| 4080 | B31-5A17 (Purser and Norris, 2000), + pTM61-*kan,gfp,pncA*,$vlsE_{A3}$ | lp25, lp28 | (Tan et al., 2022) |
| 4036 | 5A17 (Purser and Norris, 2000) *Δbbk32::strep* + pTM61-*kan,gfp,pncA* | lp25, lp28, cp9 | (Tan et al., 2022) |
| 4043 | 5A17 (Purser and Norris, 2000) + pTM61-*kan,gfp*,*pncA* | lp25, lp28, cp9 | (Tan et al., 2022) |
| 4446 | B31 5A4 pTM61*kan,gfp* | | (Tan et al., 2023) |
| 4452 | B31-A3 (Elias et al., 2002) *ΔospC::kan* + pTM61*gent,gfp*,$ospC_{B31-ECM}$ | cp9 | (Lin et al., 2020) |
| 4458 | B31-A3 (Elias et al., 2002) Δ*ospC::kan* + pTM61*gent,gfp*,$ospC_{B31}$ | lp28-4, lp56, cp9 | (Lin et al., 2020) |
| 4517 | 5A17 (Purser and Norris, 2000) *Δbbk32*::*strep* + pTM61*kan*,*gfp*,*pncA*,$vlsE_{A3}$ | lp25, lp28, cp9 | (Tan et al., 2022) |

**Table S2. Bacterial strains used in this work.**

| INHIBITOR | TARGET/PATHWAY | DOSES TESTED | TRANSMIGRATION | METABOLIC ACTIVITY IN CELLS | PRESENCE OF MONOLAYER INTEGRITY | % CELL INFECTED | *B. burgdorferi* LOAD |
|---|---|---|---|---|---|---|---|
| **LY294002** | PI3K (Chaussade et al., 2007). | 10-40 μM | Apparent increase* | Decreased | No | NA | NA |
| **U0126** | MEK1/2 | 10-30 μM | Apparent increase* | Decreased | No | NA | NA |
| **Cilengitide** | Integrins (Hariharan et al., 2007). | 1-10 μM | Apparent increase* | NA | No | NA | NA |
| **Filipin III** | Flotillin-dependent Caveolin-dependent | 0.83-5 μM | Apparent increase* | NA | Yes | NA | NA |
| **Dynasore** | Dynamin inhibitor of GTPase activity of dynamin 1/2 (it inhibits dynamine mediated endocytosis) (Macia et al., 2006). | 80 μM | No change | NA | NA | NA | NA |
| **Dasatinib** | Src, Abl (Shah et al., 2006). | 1-20 nM | No change | NA | NA | NA | NA |
| **CK-666** | Arp2/3 complex cell-permeable inhibitor (Nolen et al., 2009). | 4-30 μM | No change | Decreased | Yes | NA | NA |
| **Imipramine** | EGFR/PKC-δ/NF-κB signaling. Macropinocytosis inhibitor (Lin et al., 2018). | 5-40 μM | No change | No changed up to 5 μM; decreased at higher doses | Yes | NA | NA |
| **SB203580** | P38 inhibitor | 10-40 μM | No change | Decreased | NA | NA | NA |
| **amiloride** | Macropinocytosis inhibitor (inhibits recruitment of Cdc42/Rac1 to the membrane (Koivusalo et al., 2010). | 6-7.5 μM | DECREASED | No changed | Yes | DECREASED | DECREASED |
| **ML141** | Cdc42 selective allosteric, reversible inhibitor. At dose higher than 100 μM, it can inhibit other Rho family GTPases such as Rac1 (Hong et al., 2013). | 10-30 μM | DECREASED | No changed | Yes | DECREASED | DECREASED |
| **EHop-016** | Rac1/3 until 5 μM; Rac1/3 and partial inhibition of Cdc42 at higher dose (Montalvo-Ortiz et al., 2012). | 0.1-10 μM | DECREASED at 5 μM No change when the monolayer integrity was lost (Bb can probably use a paracellular route in this case). | Decreased | Yes, up to 5 μM. Integrity lost with higher dose (10 μM). | INCREASED | INCREASED |
| **NSC 23766** | Inhibitor of Rac1 activation by Rac- GEFs.1 (Gao et al., 2004). | 15-50 μM | DECREASED | No changed | Yes | No change | INCREASED |

**Table S3. Inhibitors tested in this manuscript: Effect on *B. burgdorferi* transendothelial migration in Transwells.**

Note: (NA) not assayed. Highlighted areas indicate relevant changes.

*Apparent increase: may be due to monolayer integrity loss.

#Median fluorescence intensity per cell (see **Figs. 8-10**).

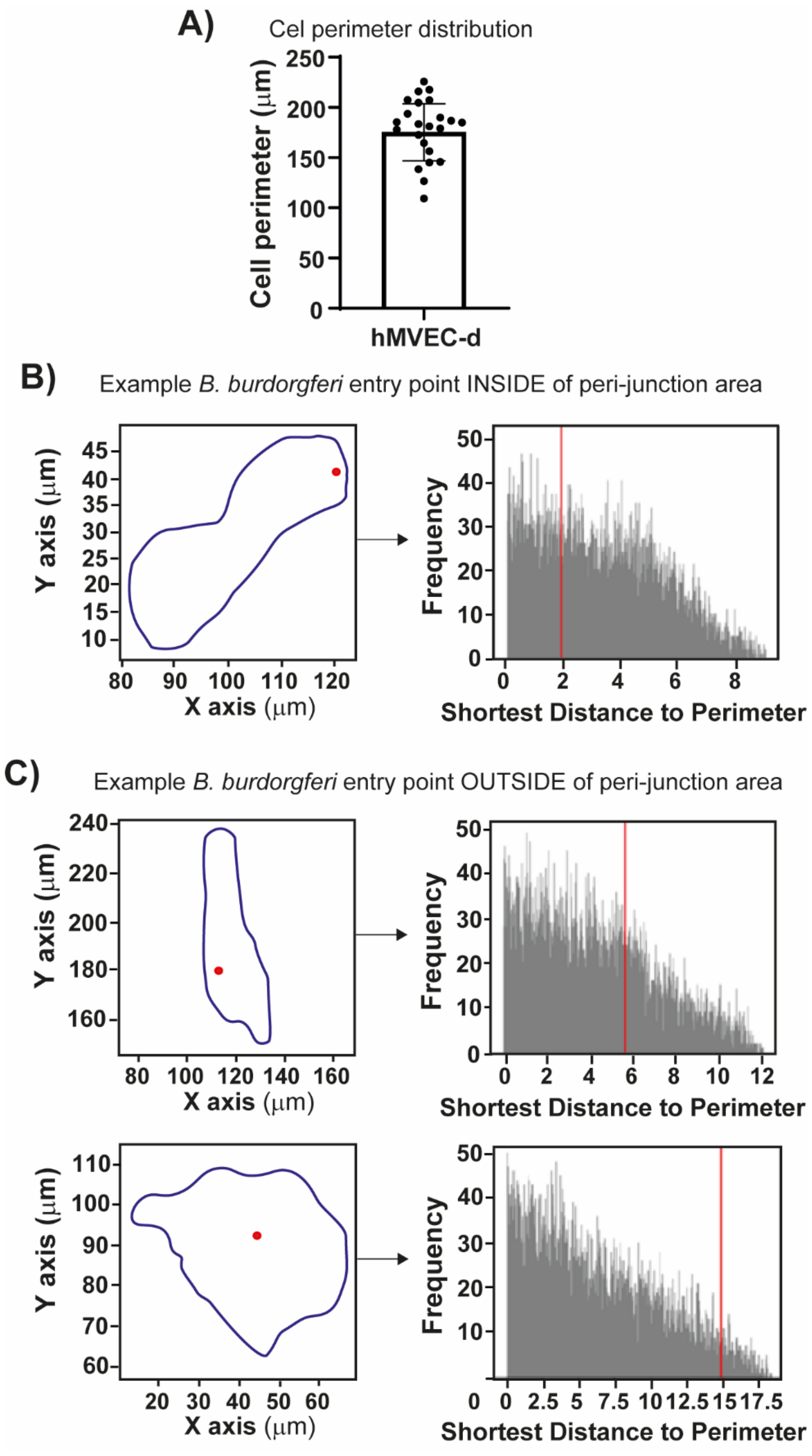
A)
Cel perimeter distribution
Cell perimeter (μm)
250
200
150
100
50
0
hMVEC-d
B)
Example B. burdorgferi entry point INSIDE of peri-junction area
Y axis (μm)
X axis (μm)
Frequency
Shortest Distance to Perimeter
C)
Example B. burdorgferi entry point OUTSIDE of peri-junction area
Y axis (μm)
X axis (μm)
Frequency
Shortest Distance to Perimeter

**Fig. S1. Examples of cell shape and B. burgdorferi entry point distribution**

**A)** Estimation of the size of the hMVEC-d cells: the perimeter of 30 cells were measured in Fiji, the average of the perimeter was 175.23 um +/- 28.36. **B)** Example of *B. burgdorferi* entry point distribution inside what was considered the peri-junction area: the micrographs showing penetrating spirochetes were used to identify coordinates of the perimeter (black) and penetration point (red). The perimeter was used to generate randomly distributed points from a continuous uniform distribution (righ panel). Finally, the shortest distance to the perimeter was computed for the penetration point (red) and the null distribution (grey). **C)** Examples of *B. burgdorferi* entry point distribution outside of what was considered the peri-junction area. It can be appreciated the difference in shape between the cells (top and bottom panel).

A)

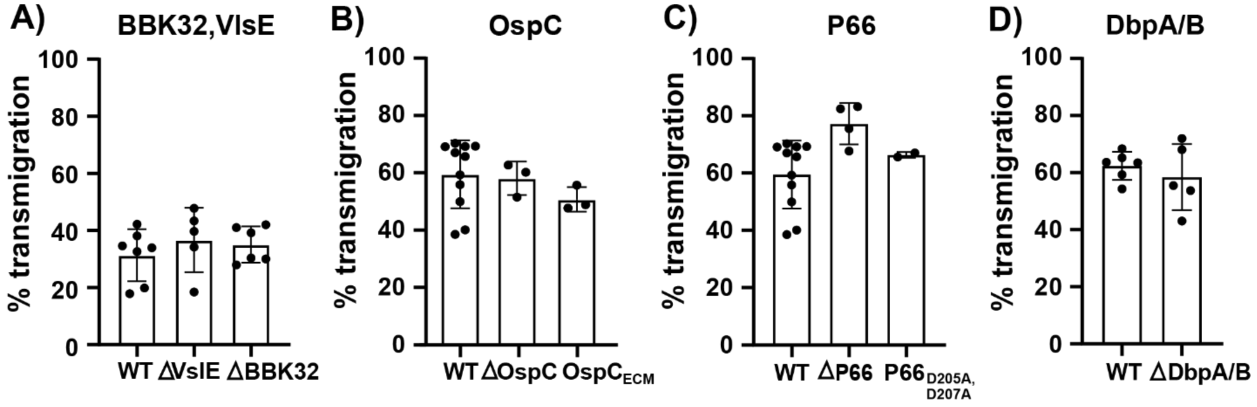


**Fig. S2. Assessment of adhesin mutants in *B. burgdorferi* transmigration. A)** Evaluation of BBK32 and VlsE: Transwell chambers were seeded with hTERT as described previously, and the upper chamber was infected with $3x10^5$ spirochetes (WT, GCB4080; VlsE knockout, GCB4043; and BBK32 knockout, GCB4017. The percentage of total transmigrated spirochetes was assessed after 20 h as described earlier. In all cases, the graphs show the percentage (mean +/- SD) of *B. burgdorferi* traversing human microvascular endothelial cells in three independent experiments performed in quadruplicate. **B)** OspC **C)** P66 **D)** DbpA/B.

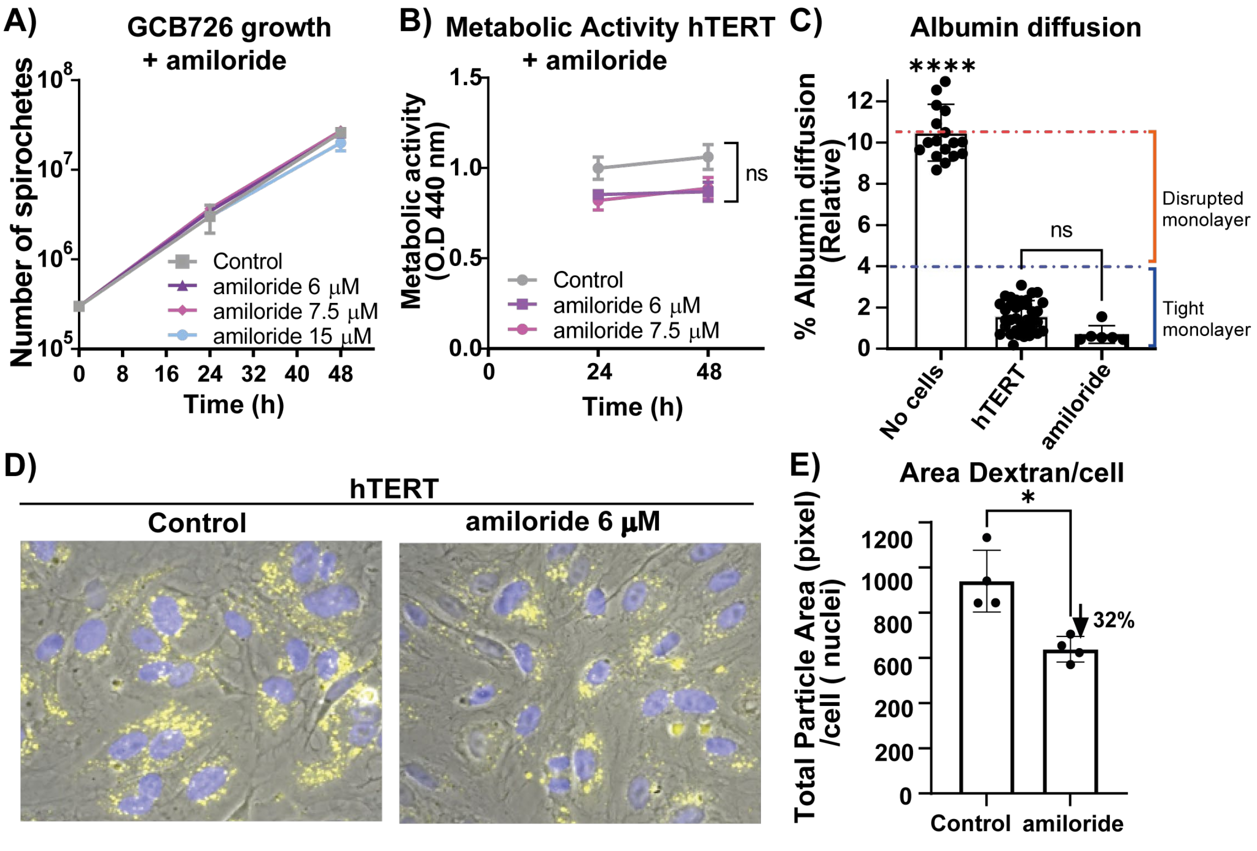


**Figure S3: Analysis of amiloride effect in B. burgdorferi and hTERT cells.**

A) Evaluation of the effect of amiloride on B. burgdorferi growth. BSK-II media lacking, or with the indicated concentrations of amiloride, was inoculated to a density of $3x10^5$ Spirochetes/ml. Spirochetes were enumerated every 24 h using a Petroff-Hausser chamber. The graph represents at least 3 independent experiments. Non-parametric ANOVA was performed to compare control versus treated cells using the Kruskal-Wallis post-test and $p < 0.05$ was considered significant. B) The metabolic activity of hTERT cells was evaluated by measuring the metabolic activity with the WTS reagent in the presence or absence of the indicated concentrations of amiloride. The cells were seeded in 96 well plates and grown to confluence in VCBM medium, then exposed to amiloride from 24-48 h. The graph represents the results of 3 independent experiments (n >10) analyzed with one-way ANOVA followed by the Kruskal-Wallis post-test. C) Evaluation of the effects of amiloride on monolayer permeability: The assessment of monolayer integrity was performed as indicated previously: 10 µg of Alexa Fluor 555-albumin was added to the upper chamber at 16 h and fluorescence was measure at 20 hours. The graph represents the mean +/- SD of three experiments performed in triplicate, measured in duplicate and analyzed with one-way ANOVA followed by the Kruskal-Wallis post-test. A single * indicates $p < 0.0332$; ** indicates $p < 0.0021$; *** indicates $p < 0.0002$; **** indicates $p < 0.0001$. D) Representative images obtained from an in vitro macropinocytosis assay. hTERT cells were grown on round coverslips to confluence and then treated with fixable-TMR-dextran or amiloride for 1h 30 min in VCBM. After fixing the cells, the nuclei were labeled with DAPI, and the cells mounted on coverslips. A phase contrast image was used to determine the area of the field covered by cells; in all cases, a confluent monolayer was analyzed for the study. Four fields of fluorescent images were captured in the rhodamine channel to show the TMR-dextran-positive macropinocytic puncta. An automatic threshold value was applied to all the images with ImageJ, macropinosomes were detected in red, and once it was accepted, the image was converted to a binary image. In those images, the macropinosomes were shown in black on a white background (Binary image), watershed processing was performed, and then the resulting image was analyzed with the particle analyzer in ImageJ. E) The graph represents the total particle area divided by the number of nuclei on each field in cells maintained in VCBM medium for 1h 30 min. More than 250 cells were analyzed. The data represent the mean +/- SD. Statistical analysis was carried out with Mann-Whitney test ($p < 0.05$) was considered significant.

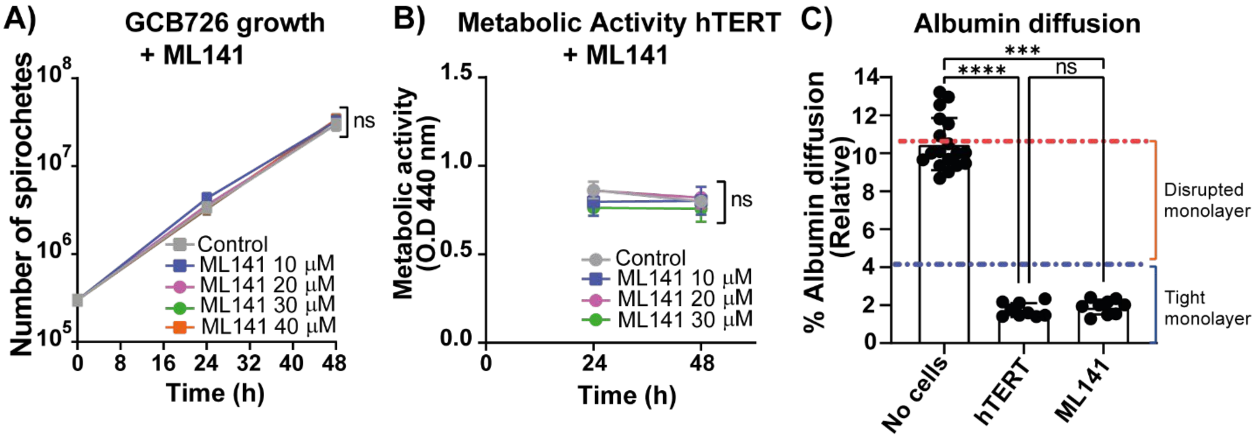


**Figure S4: Analysis of ML141 effect in *B. burgdorferi* and hTERT cells.**

**A)** Evaluation of the effect of ML141 on *B. burgdorferi* growth. BSK-II media lacking, or with the indicated concentrations of ML141 (10-40 µM), was inoculated to a density of $3X10^5$ spirochetes/ml. Spirochetes were enumerated every 24 h using a Petroff-Hausser chamber. The graph represents at least 3 independent experiments. Non-parametric ANOVA was performed to compare control versus treated cells using the Kruskal-Wallis post-test. **B)** The metabolic activity of hTERT cells was evaluated by measuring the metabolic activity with the WTS reagent in the presence or absence of the indicated concentrations of ML141 (10-30 µM). The cells were seeded in 96 well plates and grown to confluence in VCBM medium, then exposed to ML141 from 24-48 h. The graph represents the results of three independent experiments (n >10) analyzed with one-way ANOVA followed by the Kruskal-Wallis post-test. **C)** Evaluation of the effects of 30 µM ML141 on monolayer permeability. The assessment of monolayer integrity was performed as indicated previously: 10 µg of Alexa Fluor 555-albumin was added to the upper chamber at 16 h and fluorescence was measure at 20 hours. The graph represents the mean +/- SD of three experiments performed in triplicate, measured in duplicate and analyzed with one-way ANOVA followed by the Kruskal-Wallis post-test. A single * indicates $p < 0.0332$; ** indicates $p < 0.0021$; *** indicates $p < 0.0002$; **** indicates $p < 0.0001$ in all cases.

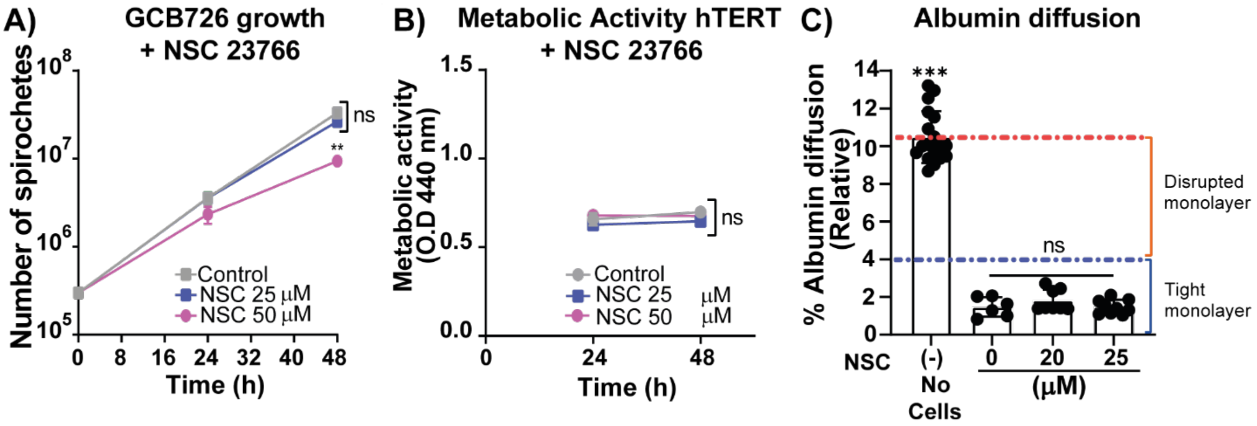


**Figure S5: Analysis of NSC 23766 effect in *B. burgdorferi* and hTERT cells.**

A) Evaluation of the effect of NSC 23766 on B. burgdorferi growth. BSK-II media lacking, or with the indicated concentrations of NSC 23766, was inoculated to a density of $3X10^5$ Spirochetes/ml. Spirochetes were enumerated every 24 h using a Petroff-Hausser chamber. The graph represents at least 3 independent experiments. Non-parametric ANOVA was performed to compare control versus treated cells using the Kruskal-Wallis post-test. B) The metabolic activity of hTERT cells was evaluated by measuring the metabolic activity with the WTS reagent in the presence or absence of the indicated concentrations of NSC 23766 (25-50 μM). The cells were seeded in 96 well plates and grown to confluence in VCBM medium, then exposed to NSC 23766 from 24-48 h. The graph represents the results of 3 independent experiments (n >10) analyzed with one-way ANOVA followed by the Kruskal-Wallis post-test. C) Evaluation of the effects of NSC 23766 on monolayer permeability: The assessment of monolayer integrity was performed as indicated previously: 10 μg of Alexa Fluor 555-albumin was added to the upper chamber at 16 h and fluorescence was measure at 20 hours. The graph represents the mean +/- SD of three experiments performed in triplicate, measured in duplicate and analyzed with one-way ANOVA followed by the Kruskal-Wallis post-test. A single * indicates $p < 0.0332$; ** indicates $p < 0.0021$; *** indicates $p < 0.0002$; **** indicates $p < 0.0001$ in all cases.

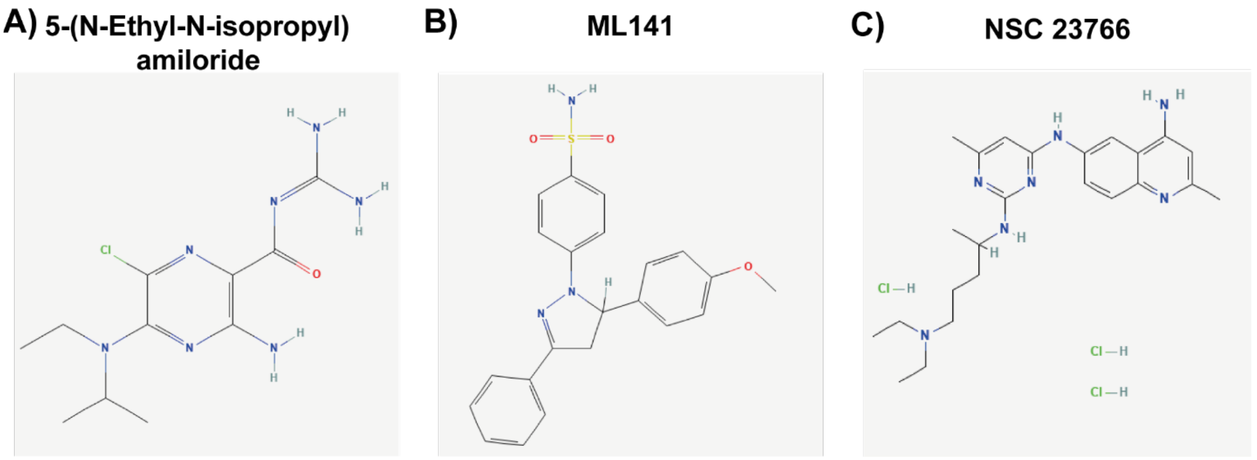

**Figure S6: Chemical Structure of the inhibitors used.**

A) 5-(N-Ethyl-N-isopropyl)amiloride, used to inhibit the membrane recruitment of Cdc42 and Rac1. B) ML141 used to selectively inhibit Cdc42. C) NSC 23766 trihydrochloride used to inhibit Rac1. Structures from PubChem.

### References


Bono, J.L., Elias, A.F., Kupko, J.J., III, Stevenson, B., Tilly, K., and Rosa, P. (2000). Efficient targeted mutagenesis in *Borrelia burgdorferi*. J Bacteriol *182*, 2445-2452.

Brissette, C.A., Kees, E.D., Burke, M.M., Gaultney, R.A., Floden, A.M., and Watt, J.A. (2013). The multifaceted responses of primary human astrocytes and brain microvascular endothelial cells to the Lyme disease spirochete, Borrelia burgdorferi. ASN neuro *5*, AN20130010.

Burns, M.J., and Furie, M.B. (1998). *Borrelia burgdorferi* and interleukin-1 promote the transendothelial migration of monocytes in vitro by different mechanisms. Infect Immun *66*, 4875-4883.

Burns, M.J., Sellati, T.J., Teng, E.I., and Furie, M. (1997). Production of interleukin-8 (IL-8) by cultured endothelial cells in response to *Borrelia burgdorferi* occurs independently of secreted IL-1 and tumor necrosis factor alpha and is required for subsequent transendothelial migration of neutrophils. Infect Immun *65*, 1217-1222.

Chaussade, C., Rewcastle, G.W., Kendall, J.D., Denny, W.A., Cho, K., Grønning, L.M., Chong, M.L., Anagnostou, S.H., Jackson, S.P., Daniele, N.*, et al.* (2007). Evidence for functional redundancy of class IA PI3K isoforms in insulin signalling. Biochem J *404*, 449-458.

Chung, Y., Zhang, N., and Wooten, R.M. (2013). Borrelia burgdorferi elicited-IL-10 suppresses the production of inflammatory mediators, phagocytosis, and expression of co-stimulatory receptors by murine macrophages and/or dendritic cells. PLoS One *8*, e84980.

Coleman, J.L., Sellati, T.J., Testa, J.E., Kew, R.R., Furie, M.B., and Benach, J.L. (1995). *Borrelia burgdorferi* binds plasminogen, resulting in enhanced penetration of endothelial monolayers. Infect Immun *63*, 2478-2484.

Elias, A.F., Stewart, P.E., Grimm, D., Caimano, M.J., Eggers, C.H., Tilly, K., Bono, J.L., Akins, D.R., Radolf, J.D., Schwan, T.G.*, et al.* (2002). Clonal Polymorphism of *Borrelia burgdorferi* Strain B31 MI: Implications for Mutagenesis in an Infectious Strain Background. Infect Immun *70*, 2139-2150.

Gao, Y., Dickerson, J.B., Guo, F., Zheng, J., and Zheng, Y. (2004). Rational design and characterization of a Rac GTPase-specific small molecule inhibitor. Proceedings of the National Academy of Sciences *101*, 7618-7623.

Grab, D.J., Nyarko, E., Nikolskaia, O.V., Kim, Y.V., and Dumler, J.S. (2009). Human brain microvascular endothelial cell traversal by *Borrelia burgdorferi* requires calcium signaling. Clin Microbiol Infect *15*, 422-426.

Grab, D.J., Perides, G., Dumler, J.S., Kim, K.J., Park, J., Kim, Y.V., Nikolskaia, O., Choi, K.S., Stins, M.F., and Kim, K.S. (2005). Borrelia burgdorferi, host-derived proteases, and the blood-brain barrier. Infect Immun *73*, 1014-1022.

Hariharan, S., Gustafson, D., Holden, S., McConkey, D., Davis, D., Morrow, M., Basche, M., Gore, L., Zang, C., O'Bryant, C.L.*, et al.* (2007). Assessment of the biological and pharmacological effects of the alpha nu beta3 and alpha nu beta5 integrin receptor antagonist, cilengitide (EMD 121974), in patients with advanced solid tumors. Annals of oncology : official journal of the European Society for Medical Oncology *18*, 1400-1407.

Hong, L., Kenney, S.R., Phillips, G.K., Simpson, D., Schroeder, C.E., Nöth, J., Romero, E., Swanson, S., Waller, A., Strouse, J.J.*, et al.* (2013). Characterization of a Cdc42 protein inhibitor and its use as a molecular probe. J Biol Chem *288*, 8531-8543.

Kawabata, H., Norris, S.J., and Watanabe, H. (2004). BBE02 disruption mutants of *Borrelia burgdorferi* B31 have a highly transformable, infectious phenotype. Infect Immun *72*, 7147-7154.

Koivusalo, M., Welch, C., Hayashi, H., Scott, C.C., Kim, M., Alexander, T., Touret, N., Hahn, K.M., and Grinstein, S. (2010). Amiloride inhibits macropinocytosis by lowering submembranous pH and preventing Rac1 and Cdc42 signaling. Journal of Cell Biology *188*, 547-563.

Kumar, D., Ristow, L.C., Shi, M., Mukherjee, P., Caine, J.A., Lee, W.Y., Kubes, P., Coburn, J., and Chaconas, G. (2015). Intravital Imaging of Vascular Transmigration by the Lyme Spirochete: Requirement for the Integrin Binding Residues of the *B. burgdorferi* P66 Protein. PLoS Pathog *11*, e1005333.

Lafrance, M.E., Pierce, J.V., Antonara, S., and Coburn, J. (2011). The *Borrelia burgdorferi* integrin ligand, P66, affects gene expression by human cells in culture. Infect Immun.

Lazarus, J.J., Kay, M.A., McCarter, A.L., and Wooten, R.M. (2008). Viable Borrelia burgdorferi enhances interleukin-10 production and suppresses activation of murine macrophages. Infection and immunity *76*, 1153-1162.

Lin, H.P., Singla, B., Ghoshal, P., Faulkner, J.L., Cherian-Shaw, M., O'Connor, P.M., She, J.X., Belin de Chantemele, E.J., and Csányi, G. (2018). Identification of novel macropinocytosis inhibitors using a rational screen of Food and Drug Administration-approved drugs. British journal of pharmacology *175*, 3640-3655.

Lin, Y.P., Tan, X., Caine, J.A., Castellanos, M., Chaconas, G., Coburn, J., and Leong, J.M. (2020). Strain-specific joint invasion and colonization by Lyme disease spirochetes is promoted by outer surface protein C. PLoS Pathog *16*, e1008516.

Ma, Y., Sturrock, A., and Weis, J.J. (1991). Intracellular localization of *Borrelia burgdorferi* within human endothelial cells. Infect Immun *59*, 671-678.

Macia, E., Ehrlich, M., Massol, R., Boucrot, E., Brunner, C., and Kirchhausen, T. (2006). Dynasore, a cell-permeable inhibitor of dynamin. Dev Cell *10*, 839-850.

Montalvo-Ortiz, B.L., Castillo-Pichardo, L., Hernández, E., Humphries-Bickley, T., De La Mota-Peynado, A., Cubano, L.A., Vlaar, C.P., and Dharmawardhane, S. (2012). Characterization of EHop-016, novel small molecule inhibitor of Rac GTPase. Journal of Biological Chemistry *287*, 13228-13238.

Moriarty, T.J., Norman, M.U., Colarusso, P., Bankhead, T., Kubes, P., and Chaconas, G. (2008). Real-time high resolution 3D imaging of the lyme disease spirochete adhering to and escaping from the vasculature of a living host. PLoS Pathog *4*, e1000090.

Nolen, B.J., Tomasevic, N., Russell, A., Pierce, D.W., Jia, Z., McCormick, C.D., Hartman, J., Sakowicz, R., and Pollard, T.D. (2009). Characterization of two classes of small molecule inhibitors of Arp2/3 complex. Nature *460*, 1031-1034.

Purser, J.E., and Norris, S.J. (2000). Correlation between plasmid content and infectivity in *Borrelia burgdorferi*. Proc Natl Acad Sci U S A *97*, 13865-13870.

Sellati, T.J., Abrescia, L.D., Radolf, J.D., and Furie, M.B. (1996). Outer surface lipoproteins of *Borrelia burgdorferi* activate vascular endothelium *in vitro*. Infect Immun *64*, 3180-3187.

Sellati, T.J., Burns, M.J., Ficazzola, M.A., and Furie, M.B. (1995). *Borrelia burgdorferi* upregulates expression of adhesion molecules on endothelial cells and promotes transendothelial migration of neutrophils *in vitro*. Infect Immun *63*, 4439-4447.

Shah, N.P., Lee, F.Y., Luo, R., Jiang, Y., Donker, M., and Akin, C. (2006). Dasatinib (BMS-354825) inhibits KITD816V, an imatinib-resistant activating mutation that triggers neoplastic growth in most patients with systemic mastocytosis. Blood *108*, 286-291.

Tan, X., Castellanos, M., and Chaconas, G. (2023). Choreography of Lyme Disease Spirochete Adhesins To Promote Vascular Escape. Microbiol Spectr, e0125423.

Tan, X., Lin, Y.P., Pereira, M.J., Castellanos, M., Hahn, B.L., Anderson, P., Coburn, J., Leong, J.M., and Chaconas, G. (2022). VlsE, the nexus for antigenic variation of the Lyme disease spirochete, also mediates early bacterial attachment to the host microvasculature under shear force. PLoS Pathog *18*, e1010511.

Thomas, D., and Comstock, L. (1989). Interaction of Lyme disease spirochetes with cultured eucaryotic cells. Infect Immun *57*, 1324-1326.

Wu, J., Weening, E.H., Faske, J.B., Hook, M., and Skare, J.T. (2011). Invasion of eukaryotic cells by *Borrelia burgdorferi* requires $\beta$1 integrins and Src kinase activity. Infect Immun *79*, 1338-1348.
